# Pan-cancer tumor classification by a holistic tumor microenvironment atlas

**DOI:** 10.64898/2025.12.27.696641

**Authors:** Shishang Qin, Xiao Du, Jinhu Li, Nan Jiang, Tian Diao, Yufei Bo, Qinhang Gao, Liangtao Zheng, Xinnan Ling, Qianqian Gao, Xiangjie Li, Sen Gao, Fei Tang, Wenjie Zhang, Chenwei Li, Peihong Fang, Linnan Zhu, Dongfang Wang, Zemin Zhang

**Author notes:** These authors contributed equally to this work (Shishang Qin, Xiao Du, Jinhu Li).

## Abstract

The tumor microenvironment (TME) heterogeneity presents a major bottleneck to effective cancer immunotherapies. We address this by establishing a pan-cancer tumor classification system based on the holistic TME cellular components. First, using single-cell transcriptomes from 1,271 patients across 26 cancer types, we compile a TME cellular atlas, leading to the identification of recurrent cell states and multicellular modules across cancer types. Among these, we highlight a type I interferon (IFN-I)-related multicellular module, including *IFIT1*^+^ tumor-associated macrophages, as a critical contributor to immune-activation TMEs. From a holistic perspective, pan-cancer TMEs exhibit trichotomous patterns predominantly driven by T, myeloid, and stromal compartments. Leveraging fine-grained TME cellular composition, we further stratify all tumors into 10 stable groups. These groups display varying responses to immune checkpoint blockade (ICB), and each rationally selected therapeutic strategy specifically perturbs the corresponding expression signature. Our pan-cancer tumor classification scheme reveals the underlying patterns of TME heterogeneity and provides a framework for stratifying patients, guiding treatment selection, and developing novel therapeutic options.

## Introduction

Tumor heterogeneity impairs the efficacy of almost all cancer therapeutics^1^, thereby becoming a central theme of cancer research. With accumulated efforts in past decades, we are approaching a comprehensive understanding of the heterogeneity of cancer cells, including their genetic, epigenetic, and transcriptional variations^2^. However, tumors are not equal to a collection of cancer cells but comprise complex ecosystems—the tumor microenvironment (TME), populated by a wide spectrum of cell types^3^. Importantly, apart from cancer cells, substantial heterogeneity manifests among the TMEs across patients and cancer types^4,5^. This TME variability profoundly influences immunotherapy outcomes. Gaining a comprehensive understanding of the TME heterogeneity can thus help identify pertinent patients who can benefit from immunotherapies and enlighten us on alternative therapeutics for other patients.

Single-cell RNA-seq (scRNA-seq) enables precise characterization of TME heterogeneity, and recent studies have explored tumor classification using single-cell-resolved TME features^6,7^. Previously, we and others have performed pan-cancer analyses of immune and non-immune cell types, revealing their widespread yet distinct differences across cancer types^7–15^. However, with a specific focus on individual or several cell types, these studies cannot capture the TME as an integrated system, obscuring the dominant cellular components that collectively define distinct tumor phenotypes. Here, we constructed a single-cell atlas for the entire TME cells from 1,271 patients across 26 solid cancer types, enabling a systematic investigation of recurrent TME patterns from a holistic perspective. First, we identified cross-lineage *IFIT1*^+^ populations and dissected their roles in the whole TME context. Second, we classified all tumors into 10 stable groups based on their holistic TME cellular compositions. This classification establishes a cellular basis for variations in immunotherapy responses and informs tailored therapeutic strategies for different patient groups.

## Results

### Construction of pan-cancer single-cell transcriptome atlas for the TME

We initiated the construction of a pan-cancer single-cell TME atlas by retrieving scRNA-seq datasets from human cancer studies. To avoid potential platform-associated variations in capturing distinct cell types, we selectively retained datasets generated by the 10x single-cell platform with unbiased sorting strategies, due to lower variations observed within the same platform (**Fig. S1a**). In total, we collected 94 scRNA-seq datasets spanning 26 solid cancer types, which were further divided into a core collection (n=55) for establishing the reference atlas and a validation set (n=39) to assess external applicability (**Fig. 1a**). We kept all treatment-naïve primary tumor samples, as well as adjacent non-tumor tissue and blood samples. After quality control, our core collection comprised a total of 4,590,413 high-quality cells from 1,192 samples of 819 patients, representing 24 cancer types (**Fig. 1b**).

**Fig. 1.**
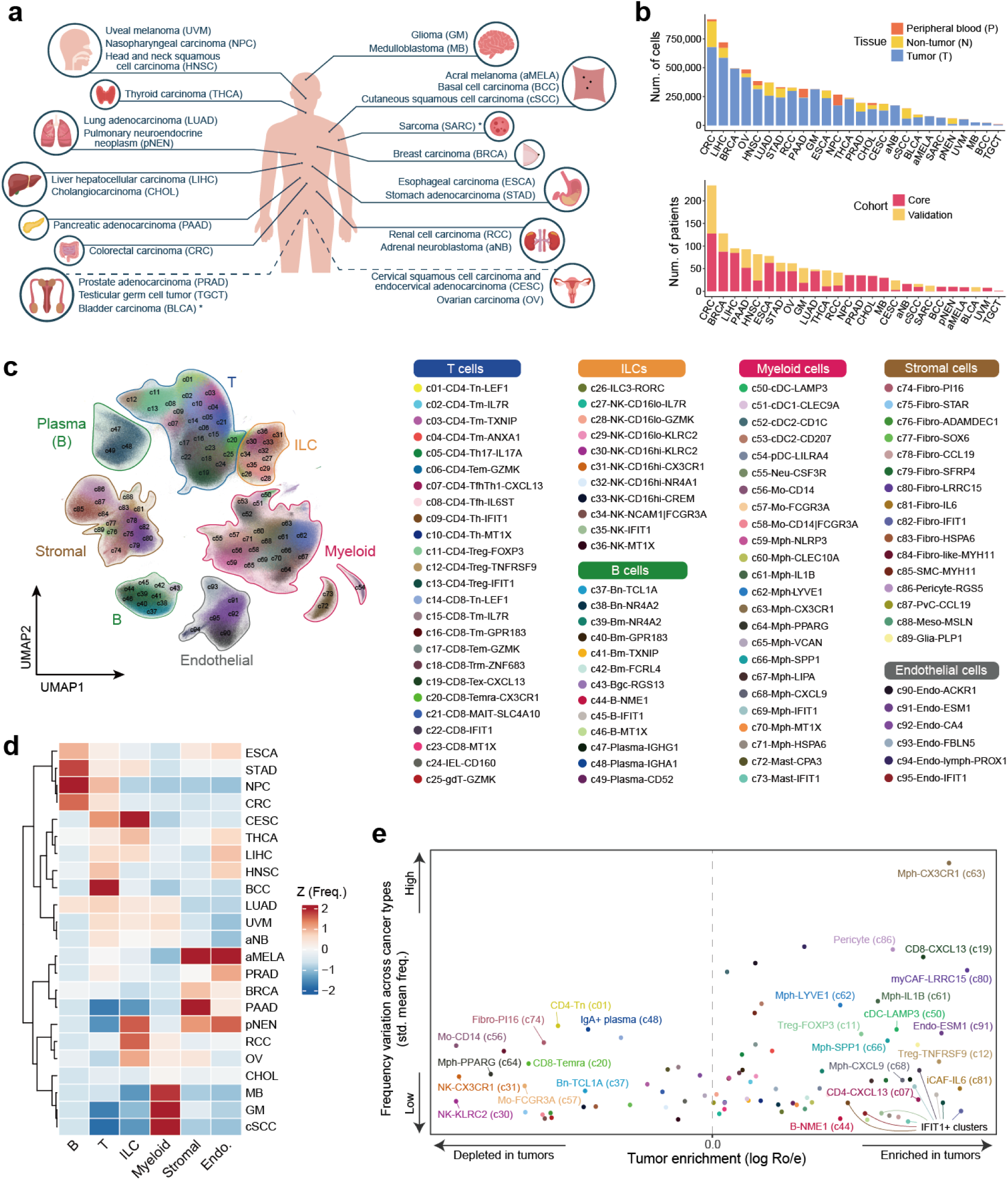
Pan-cancer single-cell transcriptome atlas for the TME. **a,** Schematic overview of cancer types involved in this study. **b,** Bar plots showing the number of cells (top) and patients (bottom) for each analyzed cancer type. **c,** Uniform manifold approximation and projection (UMAP) visualization of major cell compartments and sub-clusters. Dots represent individual cells, and colors represent different sub-clusters. **d,** Heatmap showing the relative abundance of six major compartments across different cancer types. Only cancer types from the core cohort with more than three patients are included. Color indicates the column-wise z-score-scaled median frequencies. Hierarchical clustering based on Euclidean distance is applied to rows. **e,** Scatter plot showing the tumor enrichment (x-axis) and the frequency variation across cancer types (y-axis) for each sub-cluster. Dots represent individual sub-clusters.

Integrating such an extensive collection of diverse datasets remains a computational challenge. An initial low-dimensional projection revealed that epithelial/malignant cells tended to form dataset- or patient-specific clusters, whereas immune, stromal, and endothelial cells showed a preference for aggregation (**Fig. S1b**). Accordingly, we performed malignant cell identification for each patient separately, while integrating all TME cells across datasets into a unified atlas (**Fig. S2 and Methods**). To optimize the integration of TME cells, we benchmarked several established batch-effect removal algorithms via the scIB pipeline^16^, with Harmony^17^ demonstrating superior performance and being selected for our study (**Fig. S3a**). After integration, unsupervised clustering combined with canonical marker-based annotation assigned all cells into six major compartments (**Fig. 1c**). Further sub-clustering identified 95 fine-grained subsets with distinct expression profiles (**Fig. S3b-c**), which we annotated based on literature-curated markers, supported by computational predictions from mapping to established references (**Fig. S4**). We also validated that our integration achieved effective batch correction while robustly preserving biological variation (**Fig. S5a-f and Supplementary Note1**).

The large-scale integrative analysis enabled the discovery of rare cell populations, such as *MYH11*^+^ fibroblast-like cells (c84) previously reported in lung studies^18,19^, and two *CCL19*^+^ stromal populations (c78 and c87) with distinct developmental origins^20^ (**Fig. S5g**). Notably, c78 exhibited characteristics of antigen-presenting cancer-associated fibroblasts (apCAFs)^21^ (**Fig. S5g**). To provide multifaceted characterizations of sub-clusters, we systematically evaluated their pathway activity, distribution patterns, and potential clinical associations (**Fig. S6 and Methods**). TME composition exhibited marked variations across cancer types in terms of major-compartment frequencies, while sub-clusters additionally displayed divergent patterns of tumor enrichment and cancer-type specificity (**Fig. 1d-e**). Clinical association analyses identified a set of clusters that were negatively correlated with tumor progression and positively associated with overall survival (OS) and ICB objective response rates (ICB-ORRs) (**Fig. S6 and Fig. S7a-d**). These clusters were enriched for genes involved in the activation of adaptive immune response and leukocyte adhesion (**Fig. S7d**). Conversely, six clusters linked to poor prognosis were involved in extracellular matrix (ECM) organization, vascular development, and wound healing (**Fig. S7e**), suggesting that these processes may contribute to tumor growth and immune evasion. Overall, we integrated diverse scRNA-seq datasets to build a comprehensive pan-cancer TME atlas.

### Integrative analysis of TAM heterogeneity identifies two distinct IFN-related subsets

Leveraging our atlas, we first dissected the heterogeneity of macrophages that feature highly plastic phenotypes in tumors^22^ (**Fig. 2a**). Aligning with previous observations^8^, certain resident tissue macrophages exhibited cancer-type preference, including *CX3CR1*^+^ microglia-like (c63), *PPARG*^+^ alveolar-like (c64), and *LYVE1*^+^ interstitial-like macrophages (c62) (**Fig. S8a**). We then compared the transcriptional phenotypes of distinct macrophage subtypes by examining reported functional signatures^22–26^. A recent study has proposed *SPP1*^+^ and *CXCL9*^+^ tumor-associated macrophages (TAMs) representing the macrophage polarity in tumors^27^. Consistently, we found that *SPP1*^+^ TAMs (c66) exhibited an elevated expression of multiple M2-related pathways, including response to hypoxia, ECM remodeling, and angiogenesis (**Fig. 2b**), confirming their established pro-tumoral roles. However, in addition to *CXCL9*^+^ TAMs (c68), our atlas revealed a distinct population of *IFIT1*^+^ TAMs (c69), also featuring a high expression of various M1-related pathways, such as interferon signaling, phagocytosis, and regulation of T-cell activation (**Fig. 2b**). Consistent with the pathway-level analysis, both c68 and c69 upregulated multiple T-cell recruitment and activation genes, while downregulated canonical M2 markers like *MAF* and *MRC1* (**Fig. S8b**). In addition, c68 and c69 also demonstrated distinctive transcriptional profiles and chemokine expression patterns, compared with other macrophage subsets (**Fig. S8c-d**). Notably, c69 barely expressed *CXCL9*; instead, *CXCL10* was highly expressed in both c68 and c69 (**Fig. S8c**), and achieved a high area under the curve (AUC) score when discriminating these two subsets from others (**Fig. S8e**). These two subsets constituted approximately 9% of macrophages in tumors at the pan-cancer level, with c69 relatively more abundant than c68 across most cancer types (**Fig. S8f**).

**Fig. 2.**
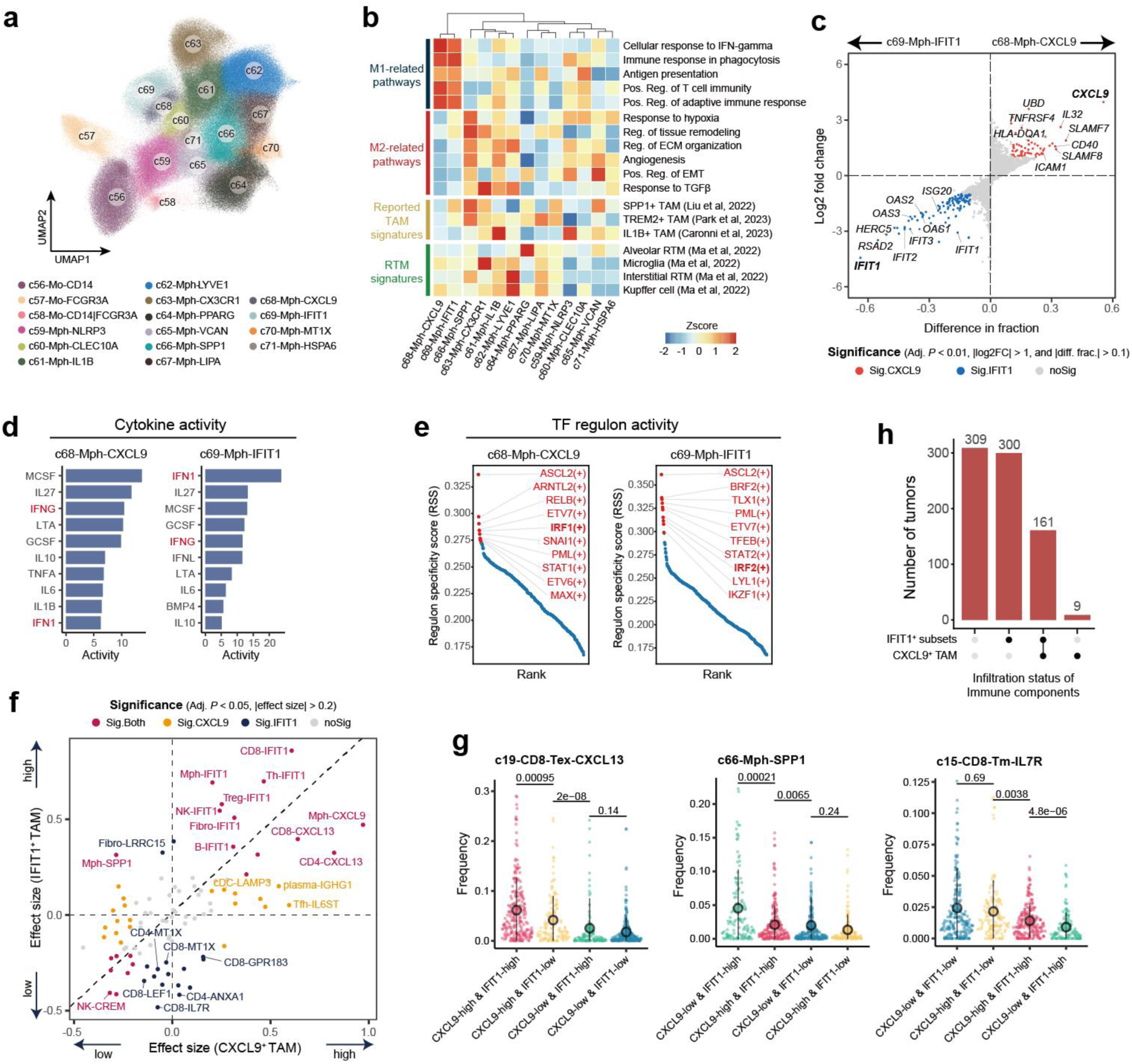
Analysis of TAM heterogeneity identifies two distinct IFN-related subsets. **a,** UMAP visualization of sub-clusters in monocytes and macrophages. Dots represent individual cells, and colors represent different cell populations. **b,** Heatmap showing the expression scores of M1- and M2-related pathways, as well as the reported TAM and RTM gene sets, in macrophage subsets. The color indicates the z-score-scaled expression of gene sets. **c,** Scatter plot showing the differential expression of genes between *CXCL9*^+^ and *IFIT1*^+^ macrophages. The x-axis represents the difference in detection fraction, while the y-axis displays log2FC. Genes with a Benjamini-Hochberg (BH)-adjusted *P* value < 0.01 (two-sided Wilcoxon test), log2FC > 1, and difference in detection fraction > 0.1 are highlighted in red, while those with log2FC < -1 are highlighted in blue. **d,** Bar plots displaying the top 10 dominant cytokines inferred by CytoSig for *CXCL9*^+^ and *IFIT1*^+^ macrophages. IFN1 and IFNG are highlighted in red. **e,** Scatter plots showing the predicted regulons for *CXCL9*^+^ and *IFIT1*^+^ macrophages, respectively. Red denotes the top regulons ranked by specificity score. **f,** Scatter plot showing effect sizes of TME cell clusters comparisons between c68-CXCL9^high^ versus c68-CXCL9^low^ and c69-IFIT1^high^ versus c69-IFIT1^low^ tumors. Dots represent cell clusters, and colors represent statistical significance categories. Effect sizes are calculated as Hedge’s g values derived from Student’s t-test, and *P* values are adjusted by the BH-method. Dots with a BH-adjusted *P* value < 0.05 and an absolute effect size >0.2 are highlighted. **g,** Sina plots comparing the frequency of representative TME cell clusters across different tumors grouped by c68-CXCL9 and c69-IFIT1. The *P* value is calculated by the two-sided Wilcoxon test. **h,** Upset plot showing the tumor numbers of different infiltration status of *IFIT1*^+^ populations and c68-Mph-CXCL9.

We next investigated differences between *CXCL9*^+^ TAMs (c68) and *IFIT1*^+^ TAMs (c69). A comparison of their expression profiles revealed that c69 exhibited elevated expression of genes and pathways related to interferon responses, particularly to type I interferons (IFN-I) (**Fig. 2c and Fig. S9a-b**). CytoSig analysis^28^ showed that c69 preferred to be regulated by IFN-I, whereas c68 exhibited high IFN-γ activity (**Fig. 2d**). We then analyzed transcription factor (TF) activity across macrophage subsets using SCENIC^29^. As expected, these two subsets shared overlapping TFs compared to other macrophages; however, two functionally antagonistic TFs, *IRF1* and *IRF2*^30^, were preferentially harbored by c68 and c69, respectively (**Fig. S2e**). These findings suggested that these two subsets might be controlled by distinct transcriptional programs.

We identified *CXCL9* (c68) and *IFIT1* (c69) as molecular markers according to their dramatic expression fold changes and differences in detection rates between these two TAM subsets (**Fig. 2c**). Notably, *in vitro* assays confirmed the specific high expression of *CXCL9* and *IFIT1* in human bone marrow-derived macrophages (BMDMs) upon the stimulation of IFN-γ and IFN-I, respectively (**Fig. S9c**). Based on these two markers, we then analyzed macrophages from a scATAC-seq dataset of human tumors^31^, revealing distinctive chromatin accessibility patterns at the *CXCL9* and *IFIT1* loci in different cell populations (**Fig. S9d**). In addition, *CXCL9*⁺ and *IFIT1*⁺ macrophages were spatially segregated across multiple tumor samples (**Fig. S9e**). Collectively, these results corroborated that *IFIT1*^+^ and *CXCL9*^+^ TAMs represented two distinct cellular states in tumors, likely induced by IFN-I and IFN-γ, respectively.

### Linking *IFIT1^+^* TAMs to the TME heterogeneity

The role of IFN-I signaling in human tumors is multifaceted, with both pro- and anti-tumor effects reported^32^. Pivoting on *IFIT1*^+^ TAMs, we systematically evaluated their associations with distinct cancer cell states and TME compositions, in comparison with *CXCL9*^+^ TAMs (**Methods**). Specifically, we divided tumors based on *CXCL9*^+^ (c68) and *IFIT1*^+^ TAM (c69) abundance, respectively, and categorized cancer cell states and TME cell clusters into a four-quadrant scheme (**Fig. 2f and Fig. S9f**).

Quadrant I tumors (c68-CXCL9^high^/c69-IFIT1^high^) harbored cancer cells with heightened interferon, cell cycle, and MYC signaling (**Fig. S9f**). Their TMEs featured abundant *CXCL13*^+^ tumor-reactive-like T cells^33,34^ and other *IFIT1*^+^ populations (**Fig. 2f-g**), hallmarks of an immune-activation TME. We performed *in vitro* IFN-I and IFN-γ stimulation to obtain macrophages resembling the phenotype of *IFIT1*^+^ and *CXCL9*^+^ TAMs and co-cultured them with T cells (**Fig. S10a-b**). These assays confirmed that these two subsets were capable of promoting T-cell activation (**Fig. S10c-d**). Additionally, we observed that *IFIT1*^+^ TAMs were more closely (above the diagonal) associated with other *IFIT1*^+^ populations (**Fig. 2f**). A recent study provides evidence that IFN-I signaling is indispensable for establishing immune-permissive TMEs, capable of triggering a robust anti-tumor T-cell response in mouse models^35^. We found that *CXCL9*^+^ TAMs rarely existed independently and were typically accompanied by at least one *IFIT1*^+^ population (**Fig. 2h**). Together with the observation that the CXCL9^high^IFIT1^high^ tumors harbored the highest *CXCL13*^+^ T cell abundance (**Fig. 2g**), we concluded that IFN-I signaling might act as a critical precondition, while the concurrently active IFN-I and IFN-γ signaling indicates a highly “hot” TME state.

Quadrant II tumors (c68-CXCL9^low^/c69-IFIT1^high^) featured a higher abundance of *SPP1*^+^ TAMs compared to other quadrants (**Fig. 2f-g**). Additionally, *LRRC15*^+^ CAFs, previously linked to the physical barrier impeding T cell infiltration^36^, were more abundant in c69-IFIT1^high^ than c69-IFIT1^low^ tumors (**Fig. 2f**). Notably, cancer cells in this quadrant upregulated epithelial-mesenchymal transition (EMT) and myogenesis signaling (**Fig. S9f**). The concurrent link to immunosuppressive populations and cancer cell plasticity suggests that IFIT1^high^ status alone does not equal to the immune “hot” phenotype.

Finally, in c69-IFIT1^low^ tumors (quadrant III/IV), cancer cells exhibited higher activity in the metal response program and peroxisome signaling (**Fig. S9f**). The TMEs of these tumors were enriched for *MT1X*^+^ populations (**Fig. 2f**), featuring signature genes that overlapped with the cancer metal response program. Metallothionein genes have been linked to cellular stress and unfavorable patient prognosis^10,14^. In addition, we found that low-activation T cells, such as CD8^+^*LEF1*^+^ and *IL7R*^+^ T cells, were overrepresented in these tumors (**Fig. 2f-g**), suggesting an immune-inactivation TME state. While the upstream drivers of these tumor characteristics remain unresolved, these analyses provided a systematic view of cancer cell and TME states linked to IFN-I signaling-related TAMs for future mechanistic investigations.

### Global cellular co-occurrence patterns in the TME

Beyond individual cell clusters, multicellular coordination underlies tissue homeostasis and disease progression^37^. To gain a holistic view of co-occurrence patterns among TME cell clusters, we constructed a cellular network based on their positive frequency correlations (**Fig. 3a**, **Fig. S11a, and Methods**). Within the network, we annotated several connected cell clusters as multicellular modules with distinct biological relevance. The robustness of these modules was supported by permutation and subsampling analyses (**Fig. S11b-c**). Furthermore, we curated 303 public Visium samples spanning 14 cancer types and mapped our fine-grained clusters onto spatial data using cell2location^38^ to evaluate the spatial colocalization tendency of these modules (**Fig. S12 and Supplementary Note2**).

**Fig. 3.**
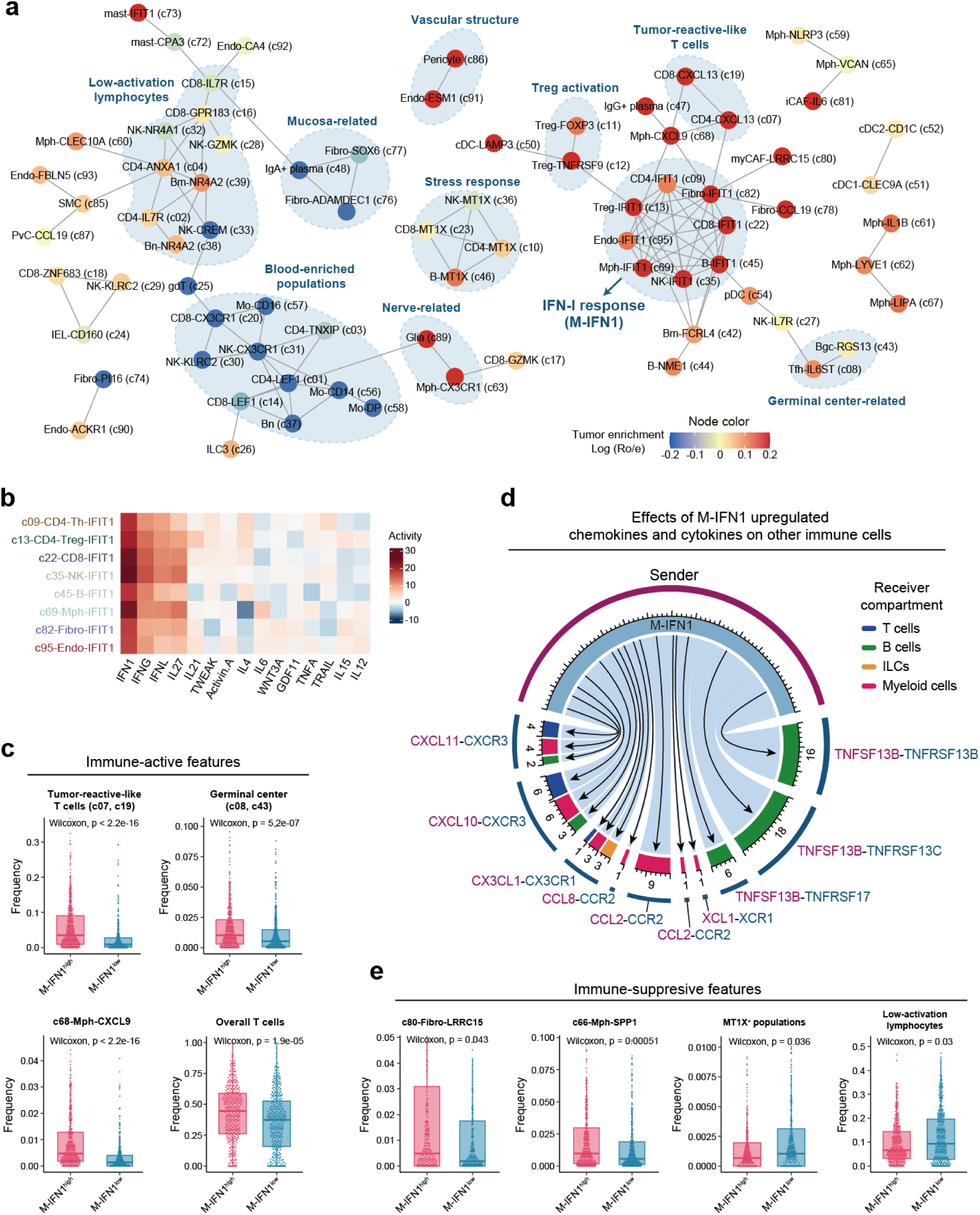
Co-occurrent cell network in tumors. **a,** Network plot showing co-occurrent cell clusters identified in tumor samples. Node colors represent the degrees of tumor enrichment of cell clusters, which are quantified by log-transformed Ro/e. Certain connected clusters with clear biological relevance are highlighted in blue. **b,** Heatmap showing the cytokine activity in M-IFN1 by the CytoSig analysis. **c,** Boxplots comparing the frequency of immune-active populations between M-IFN1^high^ and M-IFN1^low^ tumors. The *P* value is calculated by the two-sided Wilcoxon test. **d,** Circular plot showing the cell-cell interactions (CCIs) between upregulated chemokines and cytokines in M-IFN1 and other TME clusters (also see **Fig. S13d**). In the circular plot, from outer to inner, the circles represent ligand-receptor pairs and the total number of corresponding ligand-receptor pairs in each receiver compartment, respectively. Arrows from M-IFN1 point to the corresponding receiver compartment. **e,** Boxplots comparing the frequency of immune-suppressive populations between M-IFN1^high^ and M-IFN1^low^ tumors. The *P* value is calculated by the two-sided Wilcoxon test.

We found distinct biological underpinnings and spatial distribution patterns for different co-occurrent modules. First, certain modules tended to exhibit tissue-of-origin characteristics. For example, although analyzed in tumor samples, those blood-enriched clusters formed a tight module (**Fig. 3a**). The nerve and mucosa modules reflected the intrinsic organ contexts of the brain and gastrointestinal system, respectively. We noted that these two modules exhibited no spatial coherence (**Fig. S12a**), likely because their frequency correlations stemmed from co-enrichment in specific cancer types rather than spatial interactions within individual tumors (**Fig. S5f**). In contrast, cell clusters within all other annotated modules showed clear spatial co-localization (**Fig. S12a-b**). The co-occurrence of germinal center and vascular structure modules may be attributed to the close cellular communication among their clusters (**Fig. 3a and Fig. S13a**). Additionally, clusters sharing similar activation status, such as low-activation lymphocytes or high-activation *CXCL13*^+^ T cells, might co-occur in tumors (**Fig. 3a**). Finally, we detected two modules comprising clusters from different cellular compartments that potentially reacted to common stimuli—with *MT1X*^+^ populations responding to cellular stress and *IFIT1*^+^ populations to IFN-I stimulation (**Fig. 3a-b**).

Notably, the IFN-I module (M-IFN1) was highly enriched in tumors and ubiquitously detected across cancer types (**Fig. 3a and Fig. S13b**). When dividing tumors based on M-IFN1 frequencies, those with high M-IFN1 frequencies featured higher proportions of *CXCL9*^+^ TAMs, germinal center-related clusters, and overall T cells (**Fig. 3c**). These observations support the hypothesis that IFN-I stimulation is linked to the establishment of an immune-permissive condition. To gain molecular insights, we filtered ligands highly expressed by M-IFN1 populations compared to their lineage-confined counterparts and then investigated the corresponding receptors expressed by other TME populations. M-IFN1 populations, to different extents, could recruit varied immune cell types through diverse chemokine-mediated interactions (**Fig. 3d and Fig. S13c-d**). Additionally, these populations potentially supported B cell differentiation via TNFSF13B-TNFRSF13B/13C/17 axes (**Fig. 3d and Fig. S13c-d**). Collectively, M-IFN1 showed potential to facilitate the formation of an immune-hot TME. However, similar to *IFIT1*^+^ TAMs, which showed associations with immunosuppressive populations and EMT-related cancer cell states (**Fig. 2f-g and Fig. S9f**), we found that the M-IFN1^high^ tumors also harbored more *LRRC15*^+^ CAFs and *SPP1*^+^ TAMs (**Fig. 3e**). By contrast, M-IFN1^low^ tumors were enriched with low-activation lymphocytes and *MT1X^+^* populations (**Fig. 3e**), consistent with our previous findings of *IFIT1*^+^ TAMs. Together, the co-occurrence-driven analyses provided insights into multicellular organization, thereby enhancing the understanding of TME coordination.

### Tumor classification based on TME compositions

We then used our atlas to resolve the tumor heterogeneity from a holistic TME view. By interrogating all TME cellular compositions, we sought to establish a single cell-resolved, pan-cancer classification system that could comprehensively demarcate differences in the intrinsic structure of TME populations. Initial clustering based on major compartment abundances yielded three distinct groups dominated by T, myeloid, or stromal cells (**Fig. S14a**), consistent with previous findings that these lineages underlie most TME variance at the pan-cancer level^39–41^. We then refined this classification using fine-grained cell frequencies.

To determine the optimal cluster resolution, we analyzed multiple complementary statistical metrics, leading to a hyperparameter of 10 groups (**Fig. S14b and Methods**). Accordingly, we defined 10 TME subgroups (G01-G10) (**Fig. S14c**), which displayed robustness and stability under feature- and sample-based perturbations (**Fig. S14d-e and Supplementary Note3**). These subgroups broadly matched major compartment-defined groups, yet further revealed fine-grained compositional diversity (**Fig. 4a and Fig. S14f**).

**Fig. 4.**
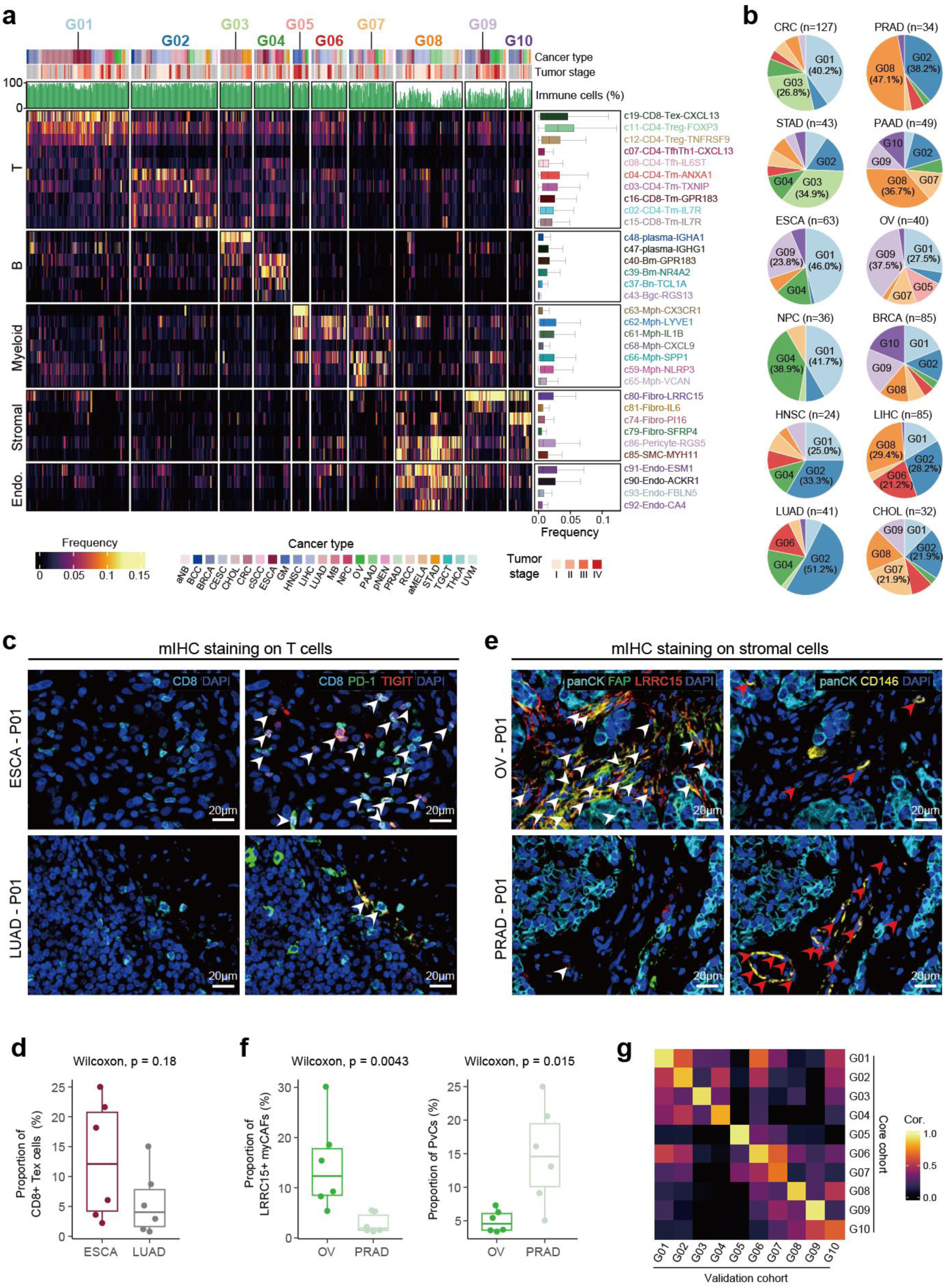
Tumor classification based on TME compositions. **a,** Heatmap showing frequencies of representative sub-clusters for 10 TME groups. For rows, box plots illustrate the distribution of sub-clusters across samples; for columns, bar plots illustrate the proportion of immune cells. **b,** Pie charts showing proportions of TME groups in different cancer types. Colors represent TME groups. Only cancer types with >20 patients are shown. **c,** Multiplex immunofluorescence staining on CD8^+^ T cells in ESCA (top) and LUAD (bottom) patients with anti-CD8 (blue), PD1 (green) and TIGIT (red) antibodies. Scale bar, 20 μm. **d,** Boxplot comparing the whole-slide proportion of CD8^+^ Tex cells between ESCA and LUAD patients, respectively. **e,** Multiplex immunofluorescence staining on stromal cells in OV (top) and PRAD (bottom) patients by anti-FAP (green), LRRC15 (red), and CD146 (yellow) antibodies. Scale bar, 20 μm. **f,** Boxplots comparing the whole-slide proportion of LRRC15^+^ myCAFs (left) and PvCs (right) between OV and PRAD patients, respectively. *P* values were calculated by two-sided Wilcoxon tests. **g,** Heatmap illustrating the Pearson correlation coefficients between the fine-grained composition of TME groups in the core set and those in the validation set.

#### G01-G04 group, T-centric tumors

G01-G04 harbored abundant T cells, corresponding to conventional immune “hot” or T-centric tumors (**Fig. S15a**). Among them, G01 uniquely enriched *CXCL13*^+^ T cells (c07 and c19) and Treg cells (c11 and c12) (**Fig. 4a and Fig. S15b**). The M-IFN1 was also prevalent in these tumors (**Fig. S15b**). Given the established correlation between CD4^+^*CXCL13*^+^ T cells and tumor mutation burden^9^, G01 might comprise tumors with high immunogenicity. In contrast, G02 exhibited enrichment of low-activation T cells despite similar overall T cell abundance (**Fig. 4a and Fig. S15b**). Additionally, cancer cells in G02 showed impaired HLA expression compared to G01 (**Fig. S16a**), indicating inferior T cell-mediated immune responses.

G03 and G04 showed few preferences for particular T cell statuses but contained abundant B cells (**Fig. S15a**). G03 exhibited high frequencies of IgA^+^ plasma cells (c48) and mucosa-related clusters (c76 and c77), primarily comprising gastrointestinal tumors (**Fig. 4a**). G04 was notable for naïve and memory B cells, along with germinal center-related clusters (**Fig. S15b**), reminiscent of tertiary lymphoid structure (TLS) characteristics^42^.

#### G05-G07 group, M-centric tumors

G05-G07 were characterized by a high proportion of myeloid cells (**Fig. S15a**), representing myeloid (M)-centric tumors. G05 harbored abundant *CX3CR1*^+^ microglia-like macrophages (c63) and mainly consisted of glioma (GM) and medulloblastoma (MB) tumors (**Fig. 4a and Fig. S15b**). Additionally, both G05 and G07 demonstrated an elevated presence of *SPP1*^+^ TAMs (c66); however, G07 also featured another angiogenesis-related *VCAN*^+^ macrophage (c65) and spanned a broad range of cancer types (**Fig. 4a and Fig. S15b**). Thus, G05 and G07 represented protumor-related M-centric groups—G05 being cancer type-specific and G07 pan-cancer-shared. By contrast, G06 exhibited high *CXCL9*^+^ macrophage (c68) prevalence alongside uniquely elevated T cell abundance among M-centric tumors (**Fig. S15a-b**), representing an M-T-centric phenotype.

#### G08-G10 group, S-centric tumors

G08-G10 exhibited low immune-cell infiltration but high proportions of stromal cells (**Fig. S15a**), indicative of immune “cold” or stroma (S)-centric tumors. Within these, G08 was distinguished by abundant endothelial cells and perivascular cells (PvCs) (**Fig. 4a and Fig. S15b**), suggesting active angiogenesis. In contrast, G09 and G10 showed higher fibroblast abundances but differed in subsets: G09 enriched tumor-specific clusters like *LRRC15*^+^ myCAFs (c80), whereas G10 favored fibroblasts prevalent in normal tissue, such as *PI16*^+^ fibroblasts (c74) (**Fig. 4a and Fig. S16b**). Additionally, G10 patients were diagnosed at earlier clinical stages than G09 (**Fig. S16c**), suggesting that these two groups represent CAF-enriched tumors at distinct progression phases.

In summary, our single-cell, holistic TME-based framework resolved both immune and non-immune high-resolution distinctions, uncovering previously unrecognized heterogeneity and advancing the understanding of pan-cancer TME diversity.

### Distribution of cancer types among TME groups

We next examined the association of TME groups with cancer types and accessible clinical parameters. Minimal associations were observed for age and sex, whereas the tumor stage exhibited modest correlations with TME groups (**Fig. S16d**). For cancer types, apart from G03 and G05, other TME groups exhibited a mixture of various cancer types (**Fig. S16e**). Our pan-cancer analysis enabled the cross-cancer type comparisons in terms of their TME group compositions (**Fig. 4b**). First, more than half of the tumors in colorectal carcinoma (CRC), esophageal carcinoma (ESCA), nasopharyngeal carcinoma (NPC), and lung adenocarcinoma (LUAD) were categorized into the T-centric G01-G04. However, they featured uneven distributions over these four groups (**Fig. 4b**). For ESCA, NPC, and CRC, a high portion of tumors were categorized into G01. In contrast, the majority of LUAD tumors were classified as G02 (**Fig. 4b**), and we confirmed that these lung tumors correspondingly harbored a low proportion of exhausted T cells (**Fig. S16f**). To further validate these findings, we analyzed six LUAD and six ESCA samples using multiplexed immunofluorescence (mIHC) staining to locate CD8^+^ Tex cells (**Fig. 4c and Fig. S17a**). The whole-slide quantification analysis revealed that LUAD samples had a lower proportion of CD8^+^ Tex cells among all CD8^+^ T cells, compared to ESCA samples (**Fig. 4d**). For pancreatic adenocarcinoma (PAAD), prostate adenocarcinoma (PRAD), and ovarian carcinoma (OV)—conventional “cold” cancer types, we observed 61%, 50%, and 43% of their tumors, respectively, belonging to the S-centric groups G08-10. In particular, PAAD and PRAD possessed a larger proportion of G08 tumors, whereas OV had more G09 tumors (**Fig. 4b**). mIHC staining on newly collected samples also confirmed the enrichment of LRRC15^+^ myCAFs and depletion of CD146^+^ PvCs in OV, in contrast to PRAD (**Fig. 4e-f and Fig. S17b**). Finally, in cancer types like LIHC (liver hepatocellular carcinoma), cholangiocarcinoma (CHOL), and breast carcinoma (BRCA), T-, M-, and S-centric groups all occupied certain proportions (**Fig. 4b**), underscoring the importance of subtyping in these cancer types. Together, these findings highlighted extensive TME heterogeneity within and across cancer types.

### Application to the large-scale validation set

To demonstrate the applicability of our classification system to new datasets, we extended our analysis to the validation set comprising 39 datasets across 16 cancer types, including bladder carcinoma (BLCA) and sarcoma (SARC) that were not present in the core collection. With the same quality control pipeline, a total of 2,171,806 high-quality cells from 672 samples of 452 patients were obtained (**Fig. 1b**). We applied TOSICA^43^, a transformer-based method, to automatically annotate cell cluster identities in the validation set using our core atlas as the reference. Notably, TOSICA outputs a probability score to quantify the prediction uncertainty. In the validation set, an averaged prediction probability of 0.94 was achieved, suggesting that for most validation cells, TOSICA could reliably assign a cell cluster label from the core atlas. Particularly, high prediction probabilities were also obtained for cells of BLCA and SARC samples (**Fig. S18a**). In addition, transcriptional profiles of predicted sub-clusters in the validation set closely mirrored their counterparts in the reference (**Fig. S18b**). These results indicated that our core atlas comprehensively covered most cell types within the TME at the pan-cancer level.

We next asked whether similar cell population structures were shared for the same cancer type between the core and validation sets. To address this, we first calculated cell subset frequencies for each cancer type in the core and validation set, observing that most cancer types tended to feature highly similar cellular compositions between the two sets (**Fig. S18c**). Notably, the characteristic sub-clusters within the 10 TME groups showed strong positive correlations between these two sets from the perspective of cancer types (**Fig. S18d**). We then trained a decision tree-based classifier on the core collection to distinguish the TME group and applied the resulting model to assign TME group labels to the validation set (**Methods**). The predicted groups in the validation set exhibited cellular composition patterns closely matching those in the reference (**Fig. 4g and Fig. S19a-b**), confirming prediction robustness. TME group distributions also showed high consistency between two sets for most cancer types (**Fig. S19c-d**), indicating stable capture of TME heterogeneity patterns for these cancer types.

### The association between TME groups and cancer cell characteristics

Although our tumor classification system was derived exclusively from TME compositions, we investigated whether certain cancer cell characteristics correlated with specific TME groups. We first analyzed transcriptional states of cancer cells, systematically comparing their expression signatures across all TME groups to detect enrichment patterns (**Methods**). As expected, G03 and G05 cancer cells highly expressed organ-specific transcriptional programs (**Fig. 5a**). The remaining groups showed diverse cancer cell state associations. For example, G01 cancer cells were linked to interferon, cell cycle, and MYC signals (**Fig. 5a**). This aligns with prior evidence that MYC dysregulation in cancer cells is related to PD-L1 upregulation and the induction of T cell exhaustion^44^. Notably, the partial EMT (pEMT) program—a hybrid epithelial-mesenchymal state^45^—was significantly enriched in G01, G04, and G09 (**Fig. 5a**). The presence of pEMT in both T- and CAF-enriched tumors extends to recent observations of its TME-dependent prognostic divergence^46^. Interestingly, G09 also harbored abundant M-IFN1 (**Fig. S15b**), further suggesting that IFN-I signaling alone does not necessarily indicate an immune-activation microenvironment.

**Fig. 5.**
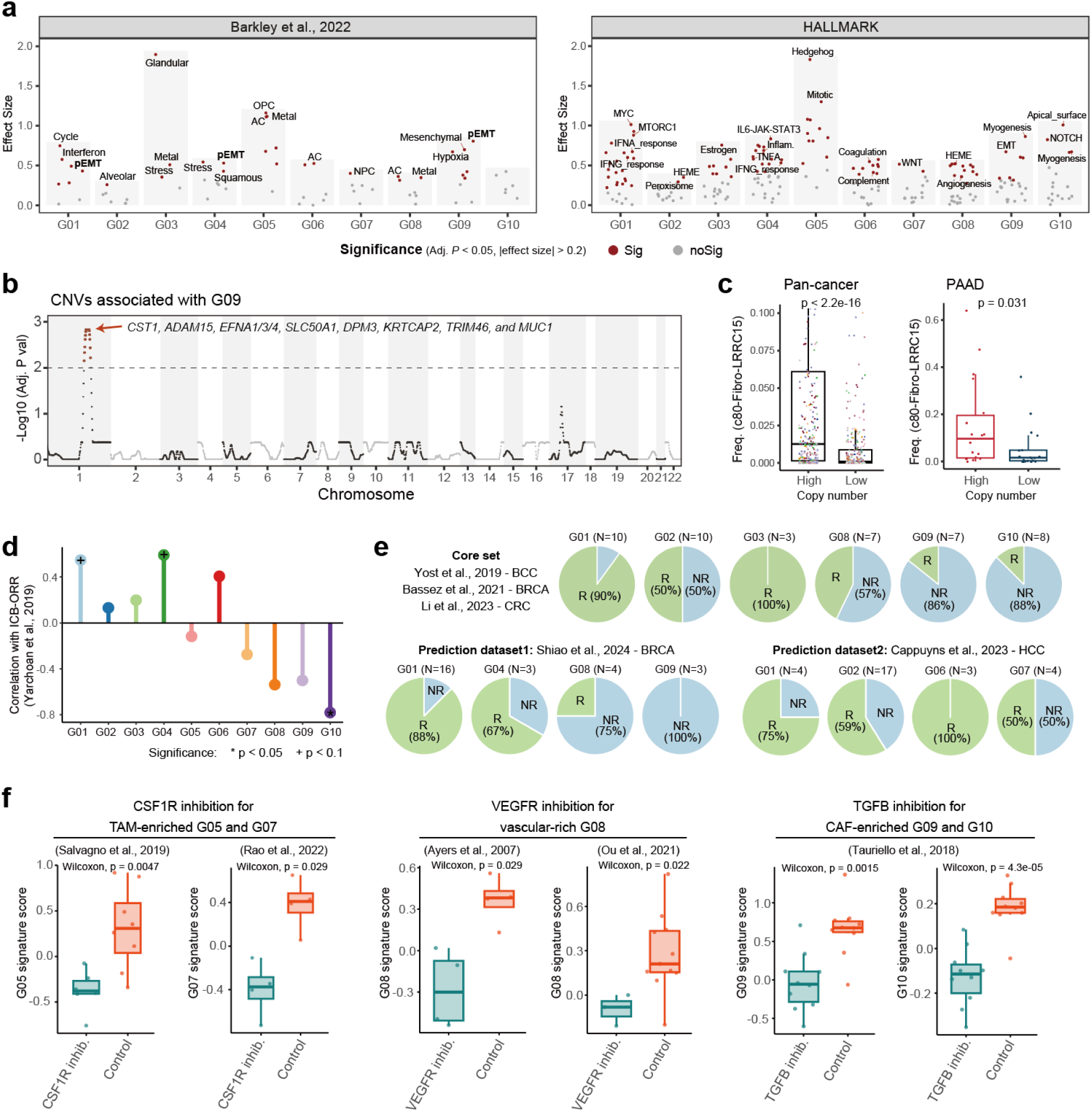
Associations of TME-based tumor classification with cancer cell states and therapeutic regimens. **a,** Scatter plots showing effect sizes of the cancer cell state comparing one TME group versus all others. Left: recurrent cancer gene programs (Barkley et al.); right: HALLMARK pathways (MsigDB). Dots represent cancer cell states, and those with a BH-adjusted *P* value < 0.05 and an effect size >0.2 are highlighted in red. Only dots with positive effect sizes are shown. Effect sizes are calculated as Hedge’s g values derived from Student’s t-test, and *P* values are adjusted by the BH-method. **b,** Scatter plot showing the significance of the associations between inferred CNVs in different chromosomal regions and the G09 group. Chromosomal regions are ordered by genomic coordinate, with alternating white and gray backgrounds distinguishing each chromosome. *P* values are derived from linear regression models and adjusted using the BH-method. The dashed line marks an adjusted *P* = 0.01. **c,** Boxplots comparing frequencies of c80-Fibro-LRRC15 across samples grouped by the CNV status (all tumors, left; PAAD only, right). The *P* value is calculated using a two-sided Wilcoxon test. **d,** Lollipop plot illustrating the Pearson correlation coefficients between the ICB objective response rate (ORR) and the proportion of each TME group within the corresponding cancer types. Only cancer types with >20 patients are included. Significant associations are labelled with ‘+’ for *P* < 0.1 and ‘*’ for *P* < 0.05. **e,** Pie charts showing the distribution of responders (R) and non-responders (NR) in each TME group using three integrated scRNA-seq datasets within our pan-cancer collection and two predicted datasets. **f,** Box plots comparing G05 and G07 signature scores between the CSF1R inhibition and control groups (left), G08 signature score between the VEGFR inhibition and control group (medium), and G09 and G10 signature scores between the TGFB inhibition and control group (right) in external expression datasets (also see **Fig. S21f**). *P* values are calculated using the two-sided Wilcoxon test.

To further dissect differences between G09 and G01/G04, we performed differential expression and CellChat-based interaction analyses. Compared to G01/G04, G09 cancer cells were enriched for multiple pathways in ECM and collagen assembly (**Fig. S20a**). Interaction analysis revealed a collagen-dominated crosstalk network in G09, with stromal cells expressing elevated *COL1A1/2* and *COL6A1/3*, mediating interactions with cancer cells (**Fig. S20b-c**). Collagen I and VI are known to promote a dense ECM that can impede T-cell infiltration^47,48^. Conversely, G09 stromal cells expressed lower levels of *COL4A1/2* (**Fig. S20d**), which encode collagen IV for basement membrane formation, and this alteration is closely linked to cancer metastasis^47,48^. Interestingly, G09 endothelial cells expressed high levels of *COL4A1/2*, potentially promoting interactions with cancer cells (**Fig. S20c-d**) and contributing to tumor angiogenesis^49^. These observations highlighted a systematic alteration of collagen expression patterns in G09 tumors, associated with an immune-cold fibrotic TME.

We next examined potential genetic cues in cancer cells, analyzing the associations between RNA-inferred CNV profiles and TME groups (**Methods**). Intriguingly, among all TME groups, G09 exhibited significant correlations with CNVs on chromosome 1q, and the peak region contained 10 genes (**Fig. 5b**). Among them, *ADAM15* and *EFNA1* are known to enhance ECM remodeling and EMT in solid tumors^48,50^. Further analysis revealed that tumors with higher CNV scores in this region harbored more *LRRC15*^+^ myCAFs across multiple cancer types (**Fig. 5c**), a finding that was independently validated in the TCGA dataset (**Fig. S20e**). Collectively, these results suggest that CNVs in cancer cells may affect the TME of G09 tumors.

### TME-based tumor classification correlated with the efficacy of therapeutic regimens

Finally, we investigated whether our proposed stratification could discern ICB responders and suggest alternative therapeutic strategies for non-responders. First, the TME group composition of cancer types was correlated with their reported ICB-ORRs^51^. As anticipated, G01 and G04 exhibited strong positive correlations with ICB response (**Fig. 5d**). Notably, a comparable correlation was observed for the M-T-centric G06 group (**Fig. 5d**). In contrast, despite harboring a high proportion of T cells, G02 displayed only a modest association with ICB-ORRs (**Fig. 5d**), aligning with the aforementioned inferior T cell-mediated responses in these tumors. These observations highlighted the indication of our TME groups in grading the treatment efficacy of tumors with abundant T cells. Conversely, *SPP1*^+^ TAMs-enriched (G05/G07) and stroma-centric groups (G08-G10) all demonstrated negative correlations with ICB-ORRs (**Fig. 5d**). To validate these associations, we analyzed three scRNA-seq datasets from our atlas and two external datasets with available follow-up ICB-response data, and observed consistent patterns (**Fig. 5e**), underscoring the response associations of our approach.

Based on the above cancer type-level ICB response analysis, we then examined gene features linked to effective ICB treatment. Using a random forest model to discriminate ICB-responsive groups (G01/G04/G06) from others, we identified a panel of 36 genes with the highest importance scores, all of which showed elevated expression in these groups (**Fig. S21a-b and Methods**). We then validated this gene panel in 11 public bulk RNA-seq cohorts spanning multiple cancer types, observing consistently higher expression of this panel in responders than non-responders (**Fig. S21c**). To benchmark predictive performance, we compared our gene panel against published signatures and the proportions of major cell types (**Methods**). Across all datasets, our panel achieved superior AUCs, demonstrating robust predictive power and broad applicability across diverse tumor contexts (**Fig. S21d**).

For G05/G07 and G08-G10 patients, representing ICB-resistant groups, we attempted to identify potential molecular regulators that could modulate the transcriptional phenotype of these TMEs. Focusing on cytokine signaling, CytoSig analysis revealed that MCSF and GCSF, both related to myeloid cell proliferation and differentiation^52^, were most active in G05/G07, while TGFB1, crucial for CAF differentiation and vascular formation^53^, dominated in G08-G10 (**Fig. S21e**). VEGFA was also highly active in G08 (**Fig. S21e**), supporting the abundant vascular-related populations observed in these tumors. Based on these observations, we posited that interventions targeting the CSF, TGFB, and VEGF pathways could potentially modulate the transcriptional states of the corresponding TME groups. To validate this, we collected public expression datasets of experiments related to these treatments and examined the treatment-mediated alterations in the signature score of each TME group. In particular, CSF1R inhibitor-treated tumor-bearing mice showed robust decreases in G05/G07 signatures relative to controls, with other TME groups remaining largely unaffected (**Fig. 5f and Fig. S21f**). Similar results were observed in VEGFR inhibition for G08 signature scores and TGFB inhibition for G08-G10 signature scores (**Fig. 5f and Fig. S21f**). These findings exemplified that tailored interventions targeting cytokines could serve as a potential strategy for modulating corresponding TMEs.

## Discussion

Tumor heterogeneity remains a roadblock to most cancer therapeutics, and decoding the underlying patterns represents a central challenge for cancer research. Here, we approached this through the lens of the holistic TME cellular components. By compiling a comprehensive single-cell transcriptome atlas, we established a pan-cancer tumor classification system with significant therapeutic implications. Our study represented a conceptual advancement that, without directly relying on cancer type-of-origins and cancer cells, TME-based classifications hold promise to tackle tumor heterogeneity and disentangle the intermingled resistant mechanisms for cancer immunotherapy.

Our comprehensive atlas provides novel insights into tumor immunity. Within macrophages, by extending the previous CXCL9-SPP1 polarization framework^27^, we identified the pan-cancer-shared *IFIT1*^+^ TAMs that represent a stable cell state critical for establishing immune-activation TMEs. Further, cross-lineage *IFIT1*^+^ populations co-occurred within tumors, with type I IFNs as common upstream regulators. Importantly, our analyses indicate a model of sequential TME activation, where IFN-I-driven signals establish a basic immune-permissive condition, and IFN-γ-driven *CXCL9*^+^ TAMs further amplify CD8⁺ T cell recruitment. This extends a recent observation in mouse tumors, in which IFN-I signaling-related monocytes are linked to anti-tumor immunity^35^. Notably, IFN-I signaling plays complex and even contradictory roles in tumor immunity^32^. We also found IFN-I signaling positively associated with immunosuppressive TME features and cancer cell plasticity. We expect future studies to elucidate the causal mechanisms underlying these observations.

Our TME-based classification system could link biological patterns to therapeutic outcomes (**Fig. S22**). To illustrate its clinical potential, we established a gene panel that encapsulated essential features of ICB responders, enabling broader identification of patients who may benefit from these therapies. For ICB-refractory groups, the holistic TME view revealed their key cellular mediators and informed tailored interventions. Importantly, previous clinical studies provide evidence to support our proposed TME modulation options. For instance, the inhibition of CSF1R and TGFB, in combination with ICB, has yielded favorable outcomes in M-centric GM and CAF-enriched PAAD, respectively^54,55^; however, the TGFB blockade in GM and TAM depletion in PAAD did not improve ICB efficacy^56,57^. For ESCA, which contains high proportions of both ICB-responsive (G01/G04) and refractory stroma-centric groups (G08-G10), a recent study reported that ICB plus TGFB inhibition achieved a high response rate of 62.5%^58^. Furthermore, ICB plus VEGFR inhibition has demonstrated effectiveness in RCC (renal cell carcinoma) and LIHC^59,60^, where vascular-rich G08 constitutes a large fraction. Collectively, these clinical studies indicated that therapies aligned with the dominant features of specific TME groups would tend to succeed, while mismatched approaches do not, further confirming the translational value of our TME classification system.

In conclusion, our study comprehensively charted the TME heterogeneity, and proposed a holistic TME-based framework for patient stratification, shedding light on the future development of tailored immunotherapy strategies.

## Methods

### Benchmark of scRNA-seq platforms

We retrieved human cells from a previous benchmark study^61^ with replicate samples sequenced by different scRNA-seq platforms. We focused our analysis on human-derived peripheral blood mononuclear cells (PBMCs), which were selected by evaluating signature genes from the original study with AUCell (v1.20.2)^29^. Cells were then annotated by our previously published SciBet algorithm (v1.0)^62^, with a reference model trained on the public pbmc3k data (https://cf.10xgenomics.com/samples/cell/pbmc3k/pbmc3k_filtered_gene_bc_matrices.tar.gz). Finally, we calculated the cellular compositions of samples under different technical conditions.

### Analysis pipeline for scRNA-seq data

#### 1) Collection and preprocessing of scRNA-seq data

We collected 94 scRNA-seq datasets that met the following criteria: i) inclusion of treatment-naïve solid cancer samples; ii) data generated by the 10x Genomics single-cell platform; iii) utilization of unbiased sorting strategies, meaning no sorting or other enrichment processes that might skew the intrinsic relative frequency of different cell types. The 94 datasets were categorized into two groups based on the publication date: the initial 55 datasets constituted our core collection, used to construct the reference atlas, while the remaining formed the validation set. Raw count matrices and corresponding detailed metadata were obtained from public databases such as Gene Expression Omnibus (GEO) or through direct contact with the original authors. Only samples from treatment-naïve primary tumors were included, along with matched adjacent non-tumor tissues and blood, the latter two being used for comparative analyses. A standardized quality control and pre-processing procedure was applied to each dataset using the scanpy pipeline (v1.9.8)^63^. Specifically, cells with total UMI counts ranging from 1,000 to 100,000, the number of detected genes ranging from 500 to 8,000, and the proportion of mitochondrial gene counts lower than 20% were retained as high-quality cells. Samples with fewer than 100 retained cells were excluded. The count matrix was normalized using the *scanpy.pp.normalize_total* function, with parameter “*target_sum=1e4*”, followed by logarithmic transformation.

We then applied a unified clustering pipeline to distinguish TME cells from malignant/epithelial cells in each dataset (**Fig. S2a**). Specifically, we excluded genes as described previously^9^ and identified highly variable genes (HVGs) across samples using the *scanpy.pp.highly_variable_genes* function, with parameter “*batch_key=sample*”. Unwanted effects, including total gene counts, the percentage of mitochondrial gene counts, and cell cycle, were regressed out using the *scanpy.pp.regress_out* function. The resulting matrix was subsequently scaled using the *scanpy.pp.scale* function. Gene blacklisting and regression were exclusively applied during unsupervised clustering to mitigate clusters arising from irrelevant or technical factors. Then, principal component analysis (PCA) was performed on the scaled matrix of HVGs using the *scanpy.tl.pca* function, and the top 50 components were retained. We next employed BBKNN (v1.5.1)^64^ to generate a batch-balanced k-nearest neighbor (KNN) graph with the sample as the batch variable. The corrected graph was then utilized to produce a Uniform Manifold Approximation and Projection (UMAP) layout using the *scanpy.tl.umap* function and to identify cell clusters using the *scanpy.tl.leiden* function with content-dependent resolution values between 0.2-0.8.

Clusters were annotated using canonical lineage markers. After annotation, TME cells proceeded to integrative clustering, while malignant/epithelial cells underwent further processing to refine malignant cell identification.

#### 2) Benchmark of batch effect correction methods

To determine the optimal batch effect removal algorithm for integrating our datasets, we employed scIB (v1.1.4)^16^ to benchmark several widely-used Python-based methods, including Harmony (v0.0.9)^17^, scVI (v1.1.2)^65^, Scanorama (v1.7.4)^66^, and BBKNN. scIB requires cell annotations for its evaluation, and we employed CellTypist (v1.6.2)^67^ with the “Immune_All_High” model to assign cell labels in a supervised manner. For computational efficiency, we conducted the benchmark analysis on a subset of 20 datasets with the lowest number of cells, including about 249k high-quality cells from 16 cancer types. Initial projection analysis indicated variances among both datasets and samples contributed to batch effects (**Fig. S1b**). Accordingly, we set the parameters for the batch removal methods as follows: i) For Scanorama and BBKNN, which support only a single batch variable, we conducted separate tests for the dataset and sample. Specifically, we configured the “*trim=30*” parameter within the BBKNN algorithm to ensure that the resulting graph remains manageable in size, thereby preventing potential memory overload^16^. ii) For scVI, which allows for multiple covariances, we set the dataset as the batch key and the sample as an additional covariate. iii) For Harmony, which also supports multiple batch variables and additionally allows for adjustable correction strength via the theta parameter, we set both the dataset and sample as batch variables and tested four combinations of theta (dataset=1 and sample=1; dataset=1 and sample=2; dataset=2 and sample=1; dataset=2 and sample=2).

We calculated 12 out of the 14 metrics provided by scIB, excluding HVG and trajectory conservation, because they did not apply to our scenarios. The overall performance scores were derived following the setting from the original publication^16^.

#### 3) Data integration and unsupervised clustering

For the core collection, the expression matrix of all datasets with epithelial/malignant cells removed was concatenated, and the resulting matrix then underwent the standardized processing steps: identifying HVGs (selecting the top 3,000 genes to accommodate the broader cancer type coverage; “*batch_key=dataset*”); regressing out the unwanted effects; scaling the matrix; and performing PCA. We then applied Harmony to correct batch effects across datasets and samples. The Harmony-corrected cellular embeddings were used to construct the KNN graph, further generating the UMAP layout and identifying Leiden-based clusters. In general, we adopted a two-round unsupervised clustering strategy to identify different cell cluster levels. In the first round of clustering, we used a resolution of 0.2 to identify clusters and annotated them as major cell types based on canonical markers. Clusters co-expressing markers from multiple lineages were considered contaminants and removed. For the T and ILC cells, an additional round of unsupervised clustering was performed to refine the distinction of CD4^+^ T cells, CD8^+^ T cells, and ILCs. Then, within each major cell type, a second round of unsupervised clustering was conducted to obtain fine-grained sub-clusters. For cluster detection, we tested resolution values ranging from 0.6 to 2.0. The optimal resolution for each second-round clustering was determined by: (i) selecting resolutions that yielded clusters with robust and biologically interpretable differential gene expression signatures; and (ii) prioritizing resolutions that produced clusters matching literature-reported cell types or states. The signature genes for each cell cluster were identified using the *scanpy.tl.rank_genes_groups* function, and those genes satisfying the following criteria were selected: i) Benjamini-Hochberg (BH) adjusted *P* value < 0.01; ii) log2FC > 1; iii) difference in detection fraction > 0.1. The sub-cluster annotations were further validated using scAnnotatR (v1.5.3)^68^ and CellTypist^67^. Specifically, for scAnnotatR, we down-sampled each annotated fine-grained cell type to 1,000 cells due to the R program’s memory constraints in large-scale single-cell data. For CellTypist, we employed the full dataset and conducted two independent validation runs using two reference models at different resolutions: ‘Immune_All_High’ and ‘Immune_All_Low’.

To tested whether our integration reliably maintained biological heterogeneity, we first assessed the preservation of global structure by comparing PCA embeddings between the uncorrected and Harmony-corrected data. For each fine-grained cell cluster, we quantified 2 metrics: intra-cluster distance (mean cosine distance from individual cells to their cluster centroid) and inter-cluster distance (mean pairwise cosine distance between cells from the two nearest neighboring clusters based on centroid proximity). Then we calculated the Pearson correlation of intra-cluster, inter-cluster distance, and their ratios between the uncorrected and Harmony-corrected data. Next, to evaluate neighborhood consistency, we identified the 15 nearest neighbors of each cell within the same sample using the *sklearn.neighbors.NearestNeighbors* function, then the Jaccard similarity was calculated between the neighbor sets in the uncorrected and Harmony-corrected data. In addition, we calculated the out-of-batch nearest neighbor (oobNN)^69^ of each cell using the *cnova.utils.calc_oobNN* function, with the *condition_key* set to the tissue-of-origins (P/N/T, defined as peripheral blood, normal tissue, or tumor tissue) of cells. Finally, we checked the distribution of oobNN in different conditions.

#### 4) Refined identification of malignant cells

To refine the identification of malignant cells from the above malignant/epithelial clusters, we utilized the recently developed Cancer-Finder software^70^ alongside the widely used inferCNV algorithm^71^. First, cells derived from tumor samples within the malignant/epithelial clusters were analyzed using Cancer-Finder (v1.0.0) with its pre-trained model, generating an initial pool of candidate malignant cells. Next, RNA-based CNV profiles of these candidate cells were inferred for each patient using a Python-optimized implementation of inferCNV (https://github.com/icbi-lab/infercnvpy, v0.5.0), with parameters “*window_size=250 step_size=10 calculate_gene_values=True*”. All normal cells from the same patient served as the reference group. Candidate malignant cells without discernible CNV patterns were excluded, and the remaining cells constituted the final refined pool of malignant cells. The resulting malignant cell identifications showed strong concordance with published DNA-based genomic studies. For example, in colorectal cancer (CRC), we observed recurrent large-fragment deletions on chromosomes 5, 14, and 18 and amplifications on chromosomes 7 and 20 (**Fig. S2b**), recapitulating prior CRC genomic hallmarks^72^. Similarly, lung adenocarcinoma (LUAD) patients exhibited frequent CNVs on chromosomes 1 and 17, while glioblastomas (GMs) displayed focal amplifications on chromosomes 7 and 10 (**Fig. S2b**), aligning with established genomic findings^73,74^. These demonstrated the robustness of our malignant cell identification pipeline.

### Systematic evaluation of cell populations

#### 1) Analysis of functional properties

To compare the functional properties of different sub-clusters in each compartment, we curated multiple gene sets based on previous studies^8–10,15,75,76^. The expression module scores were calculated using the *scanpy.tl.score_genes* function.

#### 2) Analysis of cell-cell interaction

To evaluate the cell-cell interaction (CCI) intensity of cell clusters in the TME, we performed the CellChat (v1.5.0)^77^ analysis. We subsampled the number of cells to 1,000 per sub-cluster for those clusters with more than 1,000 cells. This strategy may help minimize bias toward those with larger cell numbers, while also enhancing computational efficiency. We counted the number of predicted significant incoming and outgoing interactions for each sub-cluster.

#### 3) Tissue distribution of sub-clusters

The degree of tissue enrichment for cell clusters was quantified by comparing the observed (O) and expected (E) cell numbers, expressed as Ro/e, following our previous description^78^. The expected cell numbers were obtained from the chi-square test (*chisq.test* function). Ro/e indicated whether a certain cell sub-cluster was enriched or depleted in specific tissues. Ro/e > 1 represented a relative enrichment of the sub-cluster in a specific tissue.

#### 4) Analysis of cancer type variation

To assess the variation in cell abundance for each cell cluster, we calculated the mean frequencies among tumor samples from a specific cancer type and then computed the standard deviation of these means across different cancer types. These variance values were utilized to quantify the degree of abundance variation of sub-clusters across cancer types. A high value denotes a large variation in the frequency of this sub-cluster among different cancer types.

#### 5) Association analysis between cell populations and tumor progression

To assess the association between cell clusters and tumor progression, we applied linear models to regress tumor stages against the frequency of each cell cluster using the *lm* function in R. The cancer type was included as a covariate in these models. To ensure fair comparisons among cell clusters, min-max scaling using the function *scale_minmax* from R package dynutils (v1.0.11) was applied to normalize cell frequencies for each cluster. Then, the estimated coefficients for the tumor stage and their corresponding *P* values were used to evaluate the association degrees. The forest plots were generated using the R package forestploter (v1.1.2).

#### 6) Survival analysis of TCGA data

The TCGA Toil re-computed expression data and patient metadata were downloaded from the UCSC Xena website (https://xenabrowser.net/). Only samples derived from primary solid tumors were retained for analyses. We calculated gene scores using the *AddModuleScore* function in the Seurat package (v4.3.0)^79^, with markers of each cell cluster as input. For each cancer type, scores were separately computed and samples were stratified into high or low groups based on the median score. Then, we constructed the Cox regression models against patient overall survival with age, sex, and tumor stages as covariates, using the *coxph* function in the survival R package (v3.5.7). Finally, we combined the results of all cancer types into a pan-cancer model by the random effect meta-analysis, which was implemented by the R package meta (v7.0.0).

#### 7) Correlation of cell populations with ICB-ORRs

To investigate the potential association between cell clusters and ICB outcomes, we obtained the ICB-ORRs of multiple cancer types from a previous study^51^. Esophageal (ESCA) and gastric cancers (STAD) have been analyzed together as esophagogastric cancer in this study, while colorectal cancer has been separated into microsatellite instability/stability (CRC-MSI/MSS)^51^. Given that not all MSI/MSS information was available in our atlas, we computed the average ORR of CRC-MSI and CRC-MSS to broadly represent the overall ORR of CRC. Then, we calculated the Pearson correlation coefficient between ICB-ORRs and mean frequencies of matched cancer types in our atlas for each cell cluster. ESCA and STAD patients were merged for this analysis.

### Functional pathway enrichment analysis

We utilized the R package clusterProfiler (v4.6.2)^80^ to perform the pathway enrichment analysis. To investigate the shared functional properties of clinically relevant sub-clusters, we first conducted individual enrichment analysis on the signature genes of each sub-cluster. Then, we identified the commonly significant enriched pathways (*Q* value < 0.05) and performed the meta-analysis using the R package MetaVolcanoR (v.1.16.0).

### Assay for assessing marker specificity of *CXCL9* and *IFIT1* in macrophages

Macrophages were derived from human peripheral blood monocytes. Specifically, four milliliters of blood were obtained from a healthy donor. PBMCs were isolated using HISTOPAQUE®-1077 (Sigma-Aldrich, 10771) and centrifuged at 400g for 30 minutes at room temperature. The PBMCs were collected at the interphase and subjected to CD14^+^ separation using the EasySep™ Human Monocyte Enrichment Kit without CD16 Depletion (STEMCELL, 19058). The isolated monocytes were then seeded in a 12-well plate at a density of 1x10^6^ cells per well and cultured in Iscove’s Modified Dulbecco’s Medium (Gbico, 12440-053) supplemented with 10% fetal bovine serum (FBS) (Gbico, 10099141C) and 50ng/mL recombinant human M-CSF (ABclonal, RP01689). After a 5-day incubation, mature macrophages were obtained. Recombinant proteins IFNA1 (Novoprotein, CC75), IFNA13 (MedChemExpress, HY-P72796), IFNA14 (MedChemExpress, HY-P72243), IFNB1 (PeproTech, 300-02BC), and IFNG (PeproTech, 300-02) were then added to the medium at a final concentration of 20 ng/mL. The cells were incubated for an additional 24 hours. After incubation, all cells were collected for total RNA extraction.

Total RNA was extracted using VeZol Reagent (Vazyme, R411-01). Genomic DNA was cleared, and reverse transcription was performed using HiScript III RT SuperMix for qPCR (+gDNA wiper) (Vazyme, R323-01). Quantitative reverse transcription PCR (qRT-PCR) was conducted using ChamQ SYBR qPCR Master Mix (Vazyme, Q331-02), according to the manufacturer’s instructions. The qRT-PCR primers of genes were: *CXCL9* (Forward Primer: GTGGTGTTCTTTTCCTCTTGGG; Reverse Primer: ACAGCGACCCTTTCTCACTAC); *IFIT1* (Forward Primer: CACAAGCCATTTTCTTTGCT; Reverse Primer: ACTTGGCTGCATATCGAAAG). Data acquisition and analysis were performed on a LightCycler480 instrument (Roche).

### Assays for validating *IFIT1*^+^ and *CXCL9*^+^ macrophage phenotypes and their functions in activating T cells

OTI CD8^+^ T cells from spleens and lymph nodes of OTI mice were isolated by EasySep™ Mouse Naïve CD8^+^ T Cell Isolation Kit (Stemcell, 19858), washed and counted prior to co-culture with macrophages. Bone marrow was extracted from the femurs and tibias of C57BL/6J mice. All bone marrow cells were flushed out and filtered through a 70-µm cell strainer. After centrifugation, red blood cells were lysed. The resultant bone marrow cells were resuspended in RPMI 1640 (Gibco, 61870036) supplemented with 10% FBS (Gibco, A5669701), 1% penicillin/ streptomycin (Gibco, 15140122) in the presence of 20 ng /ml M-CSF (Abclonal, RP01216) for 5 days. For polarization, BMDMs were stimulated with 100 ng/ml of either IFNα (MCE, HY-P701053), IFNβ (MCE, HY-P73130), or IFNγ (PeproTech, 315-05) for 48 hours. The pre-stimulated BMDMs were subsequently collected for bulk RNA-sequencing analysis and co-culture with naïve OT-I CD8^+^ T cells. Then, 20,000 macrophages were plated in flat-bottom 96 well plates and incubated with 1 μg/ml SIINFEKL OVA peptide (InvivoGen, vac-sin) for 1 hour at 37°C. After two-round washes, 200,000 OTI naïve CD8^+^ T cells were added on top of the SIINFEKL pulsed macrophages at a 10:1 ratio. After 36 hours of co-culture, T cells were harvested and stained for CD8, CD44, CD25, CD62L, PD-1, CD107a to analyze their phenotypes. For intracellular cytokine detection, GolgiPlug (BD, 555029) was added to the cocultures 8 hours before the end of incubation. Intracellular staining for GZMB, IFN-γ, was performed in PermWash buffer (BD, 554723) for 1 hour. All animal experiments were approved by the Sinoresearch (Beijing)Biotechnology Co., Ltd. IACUC (approval numbers ZYZC202406021J).

### Multicellular co-occurrence analysis

To investigate cellular co-occurrence relationships in the TME, we focused on the sub-cluster pairs with positively correlated frequencies in tumor samples. We used the relative frequencies of sub-clusters within their respective major compartments to calculate the Pearson correlation coefficient for each cell type pair. Particularly, we tested a range of thresholds for eliminating low correlations, with the optimal threshold, 0.35, identified through the elbow analysis (**Fig. S11a**). The remaining highly correlated cluster pairs were formed as a cellular co-occurrent network, which was visualized by the Cytoscape software (v3.8.1)^81^. Within the network, we annotated cellular modules to demonstrate the distinct biological indications underlying the co-occurrence.

To evaluate the robustness and stability of the annotated modules, we first permuted the module memberships (keeping the number of cell clusters within each module unchanged) 1,000 times and calculated the mean correlation among cell clusters within each module for every permutation. The significance of the observed mean correlation was then determined by comparing it to the permutated null distribution. Additionally, we used the *hclust* function to conduct hierarchical clustering of cell clusters based on their Pearson correlation distance without setting any thresholds in full, 70%, 50%, and 30% subsampled data. Based on the hierarchical dendrogram, we examined whether cell clusters within the same module tended to aggregate.

For intercellular communications among particular co-occurrent cluster pairs, we obtained the statistically significant ligand-receptor pairs predicted by CellChat. To explore the impact of the M-IFN1 module on the TME, we examined significant ligand-receptor pairs between *IFIT1*^+^ populations and other TME clusters. This analysis was restricted to ligands specifically upregulated in *IFIT1*^+^ populations (**Fig. S13c**). Accordingly, we extracted the statistically significant CellChat results containing the upregulated ligands by *IFIT1*^+^ populations, while the receptors expressed by other TME clusters.

### Analyses of spatial transcriptome data

All the collected Visium datasets were first processed using the scanpy pipeline for quality control and normalization. Specifically, spots with less than 500 counts and 200 genes were excluded from downstream analysis. In addition, genes expressing in less than 10 spots were filtered out. Then the raw counts were normalized using *scanpy.pp.normalize_total* function and transformed by *scanpy.pp.log1p* function. Subsequently, we adopted cell2location (v0.1.4)^38^ to deconvolute fine-grained cell types in each spatial sample. To ensure methodological consistency and computational efficiency, we downsampled each TME cluster to 1,000 cells, establishing a unified reference for deconvolution. Following the recommended guidelines, we configured N=5 as the expected cell abundance per spot, and αy=20 to regularize within-experiment variation in RNA detection sensitivity. The output provided the estimated cell abundance for each cell type in every spatial spot.

The Xenium data obtained from the 10x Genomics website (https://www.10xgenomics.com/) was processed using Squidpy (v1.6.2)^82^. Cells with fewer than 10 total counts and genes detected in fewer than 10 cells were filtered out. After the normalization and log1p transformation, we identified cell types according to the expression of corresponding marker genes. The distribution of identified cell types was visualized by *squidpy.pl.spatial_scatter* function.

### Analyses of single-cell ATAC-seq data

Processed scATAC-seq data was sourced from Zhang et al.^31^ and the downstream analyses were conducted using Signac (v1.13.0)^83^. The peak count matrix underwent rigorous filtering to retain peaks detected in ≥ 10 cells and cells containing ≥ 200 peaks. GRCh38 gene annotations were systematically mapped to all peaks and high-quality cells were selected using multiple quality control criteria: nucleosome banding signal < 4; transcriptional start site (TSS) enrichment score > 2; fragment count in peaks between 3,000 and 20,000; ≥15% fragments in peaks; ≤5% reads overlapping genomic blacklist regions (https://github.com/Boyle-Lab/Blacklist). Macrophages were identified according to annotations from the original study. The chromatin accessibility of *CXCL9* and *IFIT1*, the marker genes specific to *CXCL9*^+^ TAMs and *IFIT1*^+^ TAMs, was validated and visualized by Signac *CoveragePlot* function.

### SCENIC analysis

Activated regulons in each macrophage subset were analyzed using the Python implementation of the SCENIC algorithm (v0.12.1)^29^. Briefly, gene-gene co-expression relationships between transcription factors (TFs) and their potential target genes were inferred using the *grn* command. Regulons, each comprising a TF and its target genes enriched for the TF’s motifs, were identified using the *ctx* command. For this step, the motif annotation database (--annotations_fname: motifs-v9-nr.hgnc-m0.001-o0.0.tbl) and the ranking databases (database_fname: hg38 refseq-r80 500bp_up_and_100bp_down_tss.mc9nr.genes_vs_motifs.rankings.feather and hg38 refseq-r80 10kb_up_and_down_tss.mc9nr.genes_vs_motifs.rankings.feather) were downloaded from the SCENIC resource repository (https://resources.aertslab.org/cistarget). Subsequently, the *aucell* command was used to calculate regulon activity for each cell. Finally, the regulon specificity scores for specific macrophage subpopulations were computed using the *regulon_specificity_scores* function from the pyscenic Python package.

### CytoSig analysis

The CytoSig can predict the signaling activity level of 43 cytokines based on transcriptome data^28^. To compare cytokine activity across different cell clusters or TME groups, we first calculated the average expression profiles for cells in each cell cluster or TME group and then predicted their respective activity scores using the *CytoSig.ridge_significance_test* function in CytoSig (v0.0.2) with parameter settings “*alpha = 1E4, nrand = 1000, alternative = ‘two-sided’* ”.

### Tumor classification

We classified all tumor samples based on their TME cellular compositional features. The Pearson correlation-based distance matrix among tumor samples was then obtained and used to perform hierarchical clustering, with the ward.D2 aggregation method. Based on the resulting dendrogram, we initially partitioned the samples into a range of groups spanning from 3 to 15. The optimal number of groups was determined by considering multiple metrics, including average distances within and between groups, internal cluster quality indices G2 and G3^84^, as well as the widely-used Davies-Bouldin index^85^ that measures the ratio of the intra-cluster distance to inter-cluster distance. Generally, these metrics indicated a candidate range of 9 to 11, and we ultimately determined 10 as the optimal number (**Fig. S13b**). The R package ComplexHeatmap (v2.14.0) was utilized for hierarchical clustering and visualization.

Robustness of the classification framework was evaluated through feature- and sample-based perturbation analyses. For feature perturbation, fine-grained cell-type fractions were perturbed by adding stochastic Gaussian noise at levels of 5%, 10%, 20%, and 30% of the original standard deviation, following a previous study^86^. Each noise level was repeated over 1,000 iterations, and the perturbed data matrices were re-clustered using the same methodology. Clustering stability was assessed using the Rand index. For sample perturbation, we conducted 1,000 iterations of bootstrapping and 50% subsampling. Clustering outcomes were also assessed using the Rand index, along with two metrics from the prior TCGA study^41^: cluster purity (CP) and normalized mutual information (NMI).

To quantify the association between TME groups and accessible clinical parameters (age, sex, cancer type, and tumor stage), we calculated the Chi-square test-based Cramer’s V index^87^. We removed pediatric MB patients with a focus on adult cancers and then divided the range of age into two groups based on the median value. We removed sex-biased cancer types, including BRCA, CESC, OV, and PRAD when calculating the relationship between TME groups with the sex.

### Differential cell communication analysis among TME groups

The differential intercellular communication analysis among TME groups was performed using the R package CellChat. Specifically, we ran CellChat for each TME group separately and merged the obtained objects using the *mergeCellChat* function. Then, we identified the upregulated genes of each TME group by the *identifyOverExpressedGenes* function and mapped the results onto the interaction network by the *netMappingDEG* function. Next, we extracted the upregulated ligand-receptor pairs in each TME group using the *subsetCommunication* function.

### Association analysis between cancer cell states and TME features

We utilized cancer gene programs defined by Barkley et al.^45^ and hallmark gene sets from MsigDB to characterize the transcriptional states of cancer cells identified in this study. To examine the association between cancer cell states and tumor groups (categorized by TAM frequencies or TME classification), we first computed the average expression profile of cancer cells within each tumor. These average profiles were then scored using the *scanpy.tl.score_genes* function with the above gene sets. For each tumor group and cancer cell state, we conducted the Student’s t-test to compare the score differences between tumors in the specific group and those in other groups. Finally, we calculated Hedge’s g values using the *esc_t* function from the R package esc (v0.5.1) to quantify effect sizes for the differences in cancer cell states.

We also investigated the relationship between cancer CNV states and tumor groups. For this analysis, CNVs calculated by inferCNV for each chromosomal region (gene bin) were utilized. Similarly, we averaged the CNV profiles of cancer cells within each tumor. Given the strong cancer-type specificity of CNV patterns, we focused on identifying recurrent associations across cancer types using a linear model with cancer type as a covariate, implemented by the *lm* function. The BH-adjusted *P* values of the model coefficients were used to screen chromosomal regions and identify CNVs significantly associated with specific TME groups.

### Multiplex immunofluorescent staining

Tumor samples from patients were fixed in 4% Paraformaldehyde for 24 hours, and then dehydrated and embedded in paraffin. Sections were stained using PanoPANEL Kits (panovue, 10234100100) according to the manufacturer’s instructions. Specifically, slides were deparaffinized with xylene and a graded series of ethanol dilutions (100%, 95%, and 70%), fixed with 10% neutral buffered formalin for 10 minutes, followed by microwave-based antigen retrieval. Antibody blocking was performed for 30 minutes. Primary antibodies were incubated for 1 hour at room temperature, and HRP-labeled secondary antibodies were incubated at room temperature for 30 minutes, followed by TSA fluorescent dye working solution incubation for 30 minutes. After multi-antigen staining, nuclei were stained with DAPI for 20 minutes. Slides were enclosed using nail polish, scanned using the SLIDEVIEW VS200 (Olympus), and analyzed with OlyVIA software. The primary antibodies-used included CD3 (1:1500, #17617-1-AP, Proteintech), CD8 (1:400, #70306, CST), PD-1 (1:300, #86163, CST), TIGIT (1:200, #99567, CST), AE1/AE3 (1:50, #M3515, Dako), FAP (1:200, #ab227703, Abcam), LRRC15 (1:200, #ab150376, Abcam), and CD146 (1:1000, #MA5-29413, Invitrogen). Secondary HRP-conjugated antibody (Cat#10013001050, Panovue) was incubated with tyramide-coupled fluorophore: Opal 780, Opal 650, Opal 570, Opal 520, and Opal 480. Stained slides were scanned with SLIDEVIEW VS200 (Olympus) and analyzed with OlyVIA analysis software (Version 3.3). The proportions of CD3^+^ CD8^+^ PD1^+^ TIGIT^+^ DAPI^+^ cells, panCK^−^ FAP^+^ LRRC15^+^ DAPI^+^ cells, and panCK^−^ CD146^+^ DAPI^+^ cells were quantified for each tumor sample in the total tumor area using HALO® platform v3.3 (Indica Labs, Albuquerque, NM, USA).

### Analyses of the validation set

The TME cells from the validation set were annotated using TOSICA^43^, a recently developed transformer-based single-cell classification method, with our core atlas as the reference. We adopted a two-round strategy for this analysis, similar to the above analyses conducted for the core set. In the first round of analysis, major cell labels were assigned to each cell, and within each major cell population, a second round of analysis was conducted to assign fine-grained sub-cluster labels. For each analysis, we independently identified 3,000 HVGs in the core atlas and used the corresponding major or fine-grained cell annotation labels for training models. The pre-prepared mask “human_gobp” was utilized during the training processes. Finally, we obtained sub-cluster labels and prediction probabilities for each cell in the validation set.

We then utilized a multi-class classifier, implemented by the *DecisionTreeClassifier* function from the Python package sklearn (v1.3.0), to map tumor samples from the validation set to our TME classification system. Specifically, we trained the model on TME groups for tumors in the core set based on their TME compositional features, and this trained model was applied to predict the TME group labels for each tumor sample in the validation set.

### Therapeutic regimens associated with TME-based tumor classification

#### 1) Validation of the association between TME groups and ICB response

To validate the associations between TME groups and ICB response in **Fig. 5d**, we collected 2 external scRNA-seq datasets, focusing exclusively on baseline, unsorted tumor samples processed via 10x Genomics to ensure consistency. Applying the same process pipeline, we assigned TME groups to individual tumor samples in these datasets and evaluated the ICB response rates for each group.

#### 2) Generating gene signatures for TME groups

To extend our analysis to other data, such as the bulk RNA-seq, we aimed to generate signature genes broadly representing our TME groups. We first constructed pseudo-bulk samples using our scRNA-seq data. Typically, scRNA-seq data may underrepresent cancer cells compared to bulk RNA-seq due to technical biases^88^. To simulate bulk-like tumor purity and account for tumor-TME composition variability, we generated pseudo-bulk profiles by computationally spiking in tumor cells at varying proportions. Specifically, we merged the normalized expression profiles of all TME cells and then added randomly selected cancer cells (with replacement) from the same sample to the merged expression profile. A series of pseudo-bulk expression profiles with tumor purity ranging from 0.3 to 0.95 (incremented by 0.05) were generated for each tumor sample. We then performed the differential gene expression analysis using the *scanpy.tl.rank_genes_groups* function from the scanpy package on this pseudo-bulk dataset. This analysis identified signature genes for each group based on the criteria of a BH-adjusted *P* value < 0.01 and log2FC > 1.

#### 3) Developing and evaluating ICB response gene panel

We developed an efficient gene panel for predicting ICB response based on our TME classification and pseudo-bulk samples. According to the cancer type-level and scRNA-seq validation analyses (**Fig. 5d-e**), G01 (Tex-enriched), G04 (TLS-enriched), and G06 (M-T-centric) were assigned as ICB-responsive groups. All others were classified as the control. Subsequently, the union of significantly upregulated genes in G01, G04, and G06 was compiled (1687 genes), forming the initial feature list for further refinement. We then employed the random forest algorithm, implemented via the sklearn package with 100 trees, to identify key genes capable of distinguishing G01, G04, and G06 from other TME groups within this pool. To mitigate potential biases stemming from unequal group sample sizes during model training, we downsampled the number of samples in each TME group to the smallest one. Using the built-in *feature_importances* function of the random forest model, we calculated the importance score for each gene. Finally, to determine the optimal gene panel, we performed elbow analysis. Specifically, we leveraged the *find_curve_elbow* function from the pathviewr package (v1.1.7) to estimate the elbow point (**Fig. S21a**). This process yielded a gene panel comprising the top 36 genes, ranked according to their importance scores.

To evaluate the association between our gene panel and ICB response, we curated 11 bulk RNA-seq datasets with annotated ICB treatment outcomes, categorizing patients as responders (R) or non-responders (NR) based on criteria defined in the original studies. Only pre-treatment samples were included in the analysis. The expression score for our gene panel was computed using the *AddModuleScore* function in the Seurat package and then min-max scaled. Subsequently, we used *glm* function to fit the ICB response and our gene panel to the logistic regression models with age, sex and tumor stage as covariates. To combine results across datasets, we applied random-effects meta-analysis using the *rma* function (metafor package v4.6-0) with restricted maximum likelihood (REML) estimation. Finally, the estimated coefficients and confidence intervals were visualized in the forest plot.

To assess the predictive performance of our gene panel, we benchmarked it against established ICB response gene signatures^89–93^ and coarse-grained cellular fractions estimated by EPIC algorithm (v1.1.7)^94^. Specifically, to enable comparability of gene expression across datasets, we performed z-score normalization within each dataset. Then, we trained an XGBoost classifier (xgboost package v1.0.0.1), incorporating age, sex, and tumor stage as covariates, with the following parameters: *objective = “binary:logistic”, eval_metric = “logloss”, max_depth = 3, eta = 0.1, nthread = 2, min_child_weight=30, lambda=2, alpha=0.5*. To balance R and NR samples, we added *scale_pos_weight* to the model. The model was trained for 10,000 iterations with early stopping after 20 rounds of non-improvement. Finally, the predictive performance was evaluated using the average area under the receiver operating characteristic curve (AUC) from 10-fold cross-validation and the AUC was calculated via the pROC package (v1.18.5).

#### 4) Analysis of transcriptome datasets related to mouse experiments

We collected post-treatment bulk RNA-seq or genome-wide microarray datasets associated with the proposed cytokine perturbation therapies. For these datasets, we compared samples from the treated group with those derived from the control group. Specifically, the comparisons were performed on the normalized and log2-transformed expression profiles. For datasets containing only raw counts, we used the *DESeq* function in the DESeq2 package (v1.34.0)^95^ for RNA-seq data normalization, and utilized the *mas5* function in the affy package (v1.72.0)^96^ for microarray data normalization. Gene scores were calculated using the *AddModuleScore* function in the Seurat package, with the top 20 signature genes specific to each TME group as inputs.

### Supplementary Note

#### Supplementary Note 1 | Assessment of integration quality and biological variation preservation during the reference atlas construction

After integration, most clusters exhibited a well-balanced mixture of diverse datasets, indicating effective integration (**Fig. S3b**). To quantitatively evaluate integration quality and assess whether the integrative clustering preserved underlying biological variation, we conducted a series of complementary analyses. Specifically, we:

- Directly compared batch-corrected and uncorrected data at both the cluster and single-cell levels to examine whether batch removal distorted meaningful biological heterogeneity (analyses #1 and #2).
- Analyzed the preservation of tissue-specific biological signals within local cell neighborhood relationships in the integrated embedding (analysis #3).
- Examined the distribution of clusters with well-characterized tissue- or organ-specificities (analysis #4).

##### 1) Preservation of distance relationships among clusters

Since our batch correction was performed in the PCA embedding space using the Harmony algorithm ^17^, all downstream clustering and visualization relied on these Harmony-corrected embeddings. Thus, we focused our comparison between the original PC space and the Harmony-corrected embedding space. Considering PCA captured dominant biological variance in scRNA-seq data, we reasoned that the biological variations could be considered retained if the Harmony space effectively preserved the original similarity/distance relationship in the PCA space.

Accordingly, we computed the intra-cluster and inter-cluster distances for each fine-grained cell type in the uncorrected and Harmony-corrected space, respectively. Specifically, the intra-cluster one was defined as the average cosine distance between each cell and the centroid of the corresponding cluster, whereas the inter-cluster one was calculated as the average distance to the nearest neighboring clusters (based on the centroid proximity). Of note, inter-cluster distances consistently exceeded intra-cluster distances in both the uncorrected and Harmony-corrected space (**Fig. S5a**). For the comparison before and after Harmony correction, we observed strong correlations for both the intra- (Pearson’s R = 0.89) and inter-cluster distances (Pearson’s R = 0.8) (**Fig. S5b**). These results validated that Harmony removes batch effects without conflating meaningful heterogeneity at the cell-cluster level.

##### 2) Local neighborhood fidelity at the cell level

To evaluate biological similarity preservation at cellular resolution, we examined the consistency of within-batch cellular neighborhoods between uncorrected (original PCA space) and batch-corrected (Harmony-integrated space) embeddings. For each cell, we retrieved its *k*-nearest neighbors (*k*=15) within its original sample in both data spaces, respectively, and calculated the Jaccard similarity index to quantify neighborhood overlap. Despite certain cluster-specific variability, all cell populations exhibited robust neighborhood preservation, with the mean Jaccard index exceeding 0.6—equivalent to over 12/15 shared neighbors per cell, for all major compartments (**Fig. S5c**). This high concordance confirmed that our integration maintained local cellular relationships.

##### 3) Preservation of tissue-specific biological signals in cell neighborhood relationships

To further validate biological fidelity in the integrated data, we employed the out-of-batch nearest neighbor (oobNN) approach, which assesses whether nearest neighbors drawn exclusively from other samples share the same biological identity^69^. For the biological condition, we chose the tissue-of-origin (peripheral blood, normal tissue, or tumor tissue) of this cell. Critically, since the sample label was involved in batch indicators, over-correction might inadvertently mix cells from different tissues. Instead, in our integration, we observed a high enrichment of same-tissue oobNNs, validating the successful preservation of tissue-specific biological signals (**Fig. S5d**).

##### 4) Preservation of tissue- or organ-specific cell clusters

Finally, we examined the distribution of clusters with well-characterized tissue- or organ-specificities. We first focused on the difference between blood samples and those from other tissues, finding that multiple populations such as naïve CD4^+^ T cells (c01) and *CD14*^+^ monocytes (c56) were abundant in blood, while few in other tissues (**Fig. S5e**), as anticipated. In addition, the previously reported organ-specific clusters were predominantly confined to their expected organs, with examples of *CD207*^+^ Langerhans-like cells (c53) in samples from the skin, *CX3CR1*^+^ microglia-like macrophages (c63) from the brain, *PPARG*^+^ alveolar-like macrophages (c64) from the lung, and *STAR*^+^ fibroblasts (c75) from the ovary (**Fig. S5f**). Gastrointestinal samples exhibited a high abundance of IgA^+^ plasma cells (c48), *ADAMDEC1*^+^ fibroblasts (c76), and *SOX6*^+^ fibroblasts (c77) (**Fig. S5f**), consistent with previous findings^97^.

Together, these multi-perspective complementary evaluations provide robust evidence that our integrative clustering approach effectively removes batch effects while preserving meaningful biological variation.

#### Supplementary Note 2 | Supporting analyses for multicellular co-occurrence modules

##### 1) Permutation testing of correlation strength

To assess the correlation strength of intra-module cell clusters, we permuted module memberships (keeping the number of cell clusters within each module unchanged) 1,000 times and calculated the mean correlation among cell clusters within each module for every permutation. The observed mean correlations were significantly higher than those derived from the permuted null distribution (*P* < 0.05) across all modules, confirming that the identified modules represent non-random and robust associations (**Fig. S11b**).

##### 2) Threshold-free hierarchical clustering and subsampling analysis

We conducted hierarchical clustering based on correlation distance without applying any thresholds. Notably, cell clusters within the same module were closely aggregated in the hierarchical clustering dendrogram (**Fig. S11c**). To further assess module stability, we performed systematic subsampling of both the core and validation datasets, retaining 70%, 50%, and 30% of samples (**Fig. S11c**). Even at 30% subsampling, the cell clusters within the same module remained tightly aggregated. These results confirm the module reliability across datasets and subsampling conditions.

##### 3) Spatial co-localization evidence of co-occurrence relationships

To evaluate the spatial colocalization tendency of the co-occurrent cellular modules, we curated 303 publicly available 10x Visium samples spanning 14 cancer types. For spatial deconvolution, we employed cell2location^38^, which has been recommended in a recent benchmark analysis^98^. To ensure methodological consistency and computational efficiency, we downsampled each TME cluster to 1,000 cells, establishing a unified reference for deconvolution. Cell2location reliably estimated the relative abundance of cell clusters across spatial spots. For example, germinal center-related clusters (Tfh and Bgc) were predominantly confined to TLS regions in the H&E images, while *CXCL13*^+^ T cells were enriched in tumor regions (representative views were shown in **Fig. S12b**).

We next sought to quantitatively evaluate the spatial colocalization tendencies of the co-occurrent cellular modules inferred from our scRNA-seq-based frequency analyses. Specifically, for each Visium slice and each module, we computed Pearson correlation coefficients for each cell type pair across all spots, and then calculated the mean pairwise correlations within the module (inner pairs) and outside the module (outer pairs). We hypothesized that inner-pair correlations would be significantly higher than outer-pair correlations for cell modules with spatial co-localization relationships. Our analyses confirmed spatial co-localization for most co-occurrence modules (**Fig. S12a**).

Specifically, we broadly categorized frequency-based co-occurrence modules into four distinct modes. Spatial analyses further delineated their specific spatial patterns.

- First, while the blood-enriched module showed certain co-localization patterns, other tissue-of-origin-associated modules (nerve- and mucosa-related) exhibited no spatial coherence (**Fig. S12a**). We reasoned that the observed frequency correlations for these two modules across samples likely stemmed from their co-enrichment in specific cancer types (e.g., mucosa-related clusters in CRC and STAD) rather than spatial interactions within individual tumors.
- The germinal center-related and vascular structure-related modules, where we inferred close cellular communications, showed robust spatial co-distribution (**Fig. S12a**). To illustrate this, we inspected the members of the germinal center-related module (Tfh and Bgc) and the vascular module (pericytes and *ESM1*^+^ endothelial cells), observing their physical proximity (**Fig. S12b**).
- Modules reflecting cellular activation states also exhibited coordinated spatial distributions (**Fig. S12a**). The module containing highly activated lymphocytes (*CXCL13*⁺ tumor-reactive-like T cells) and that of lowly activated lymphocytes (e.g., *IL7R*^+^ T cells) displayed overlapping niches of their cell type members, respectively, potentially reflecting antigen stimulation gradients (**Fig. S12b**). Similarly, the Treg module, comprising developmentally distinct subsets, showed spatially resolved coordination (**Fig. S12b**).
- Finally, modules related to cellular stress and IFN-I response, indicative of shared upstream regulation, also displayed spatial coherence (**Fig. S12a-b**).

In summary, 8/10 annotated co-occurrence modules showed harmonized spatial and frequency-based relationships, supporting their potentially coordinated roles, whereas the discrepancies in tissue-of-origin modules (nerve- and mucosa-related) exemplified how co-occurrence- and colocalization-driven analyses can provide complementary insights into multicellular organization.

#### Supplementary Note 3 | Robustness evaluation of TME classification

To evaluate the robustness and stability of our TME classification, we performed both feature- and sample-level perturbation analyses.

##### 1) Feature perturbations

We first adopted a feature perturbation-based approach from a recently published tumor classification study^86^. Specifically, fine-grained cell-type fractions were perturbed by adding stochastic Gaussian noise at four levels: 5%, 10%, 20%, and 30% of the original standard deviation, with 1,000 iterations performed for each level. The perturbed data matrices were re-clustered using the same methodology, and the resulting clusters were compared to the original ones using the Rand index as suggested by the above study. The clustering outcomes showed high stability, with a mean index value of 0.906 at 5% noise and a minor reduction to 0.885 at 30% (**Fig. S14d**), indicating that our classification framework is robust to substantial feature perturbations.

##### 2) Sample perturbations

From the sample perturbation perspective, we conducted 1,000 iterations of bootstrapping and 50% subsampling. Clustering outcomes were also evaluated using the Rand index. We regarded this as a more challenging task than the above feature perturbation one; thus, according to the previous TCGA study^41^, we included cluster purity (CP) and normalized mutual information (NMI) metrics for this analysis. Results consistently showed high stability across all perturbation strategies, with the Rand index maintaining large values (**Fig. S14e**). Despite the relatively smaller cohort size compared to the TCGA dataset, CP and NMI values remained robust, with mean and median CP and NMI values exceeding 0.6 under all conditions and narrow interquartile ranges of 0.05∼0.06 (**Fig. S14e**). These findings further support the robustness of our classification under sample variability.

Together, these perturbation analyses from both feature and sample perspectives provide strong evidence for the stability and reliability of our classification.

### Data and code availability

This study primarily utilized public datasets, with newly generated mouse RNA-seq data from experimental validation also included. The integrated single-cell data and other analyzed data, as well as analysis codes, will be made publicly accessible upon formal publication of this manuscript.

## Acknowledgements

We would like to thank members of Z.Z. laboratory for insightful discussions. This study was supported by National Key Research and Development Program of China (2023YFF1204700 and 2022YFC2505005), National Natural Science Foundation of China (62203019, 92159305, 92259205, 92459001, and L2424217). S.Q. was supported in part by the Postdoctoral Fellowship of Peking-Tsinghua Center for Life Sciences. J.L. was supported in part by the China National Postdoctoral Program for Innovative Talents (BX20250134). We thank the National Center for Protein Sciences Beijing at Peking University for assistance with microscopy experiments. We thank the Laboratory Animal Center of Peking University for advice and technical support. Part of the analysis was performed on the High-Performance Computing Platform of the Center for Life Sciences, Peking University.

## Author contributions

Z.Z., D.W., Linnan Z., and S.Q. conceived and designed the study. J.L., X.D., S.Q., Xiangjie L., and S.G. collected the data. S.Q., X.D., J.L., Qinhang G., Xinnan L., W.Z., and P.F. performed the formal analysis. N.J., T.D., Y.B., and Qianqian G. conducted the experiments. D.W., Linnan Z., Liangtao Z., and C.L. contributed to the methodology. N.J. conducted the clinical survey. F.T. and S.Q. designed the graphs. S.Q., D.W., J.L., and X.D. wrote the original draft. Z.Z., D.W., and Linnan Z. reviewed and edited the manuscript.

## Competing interests

Z.Z. is the founder of Analytical BioSciences. Liangtao Z., C.L., and P.F. are employees of Analytical BioSciences. The other authors have declared no conflicts of interest.

## Supplementary figures

**Fig. S1.**
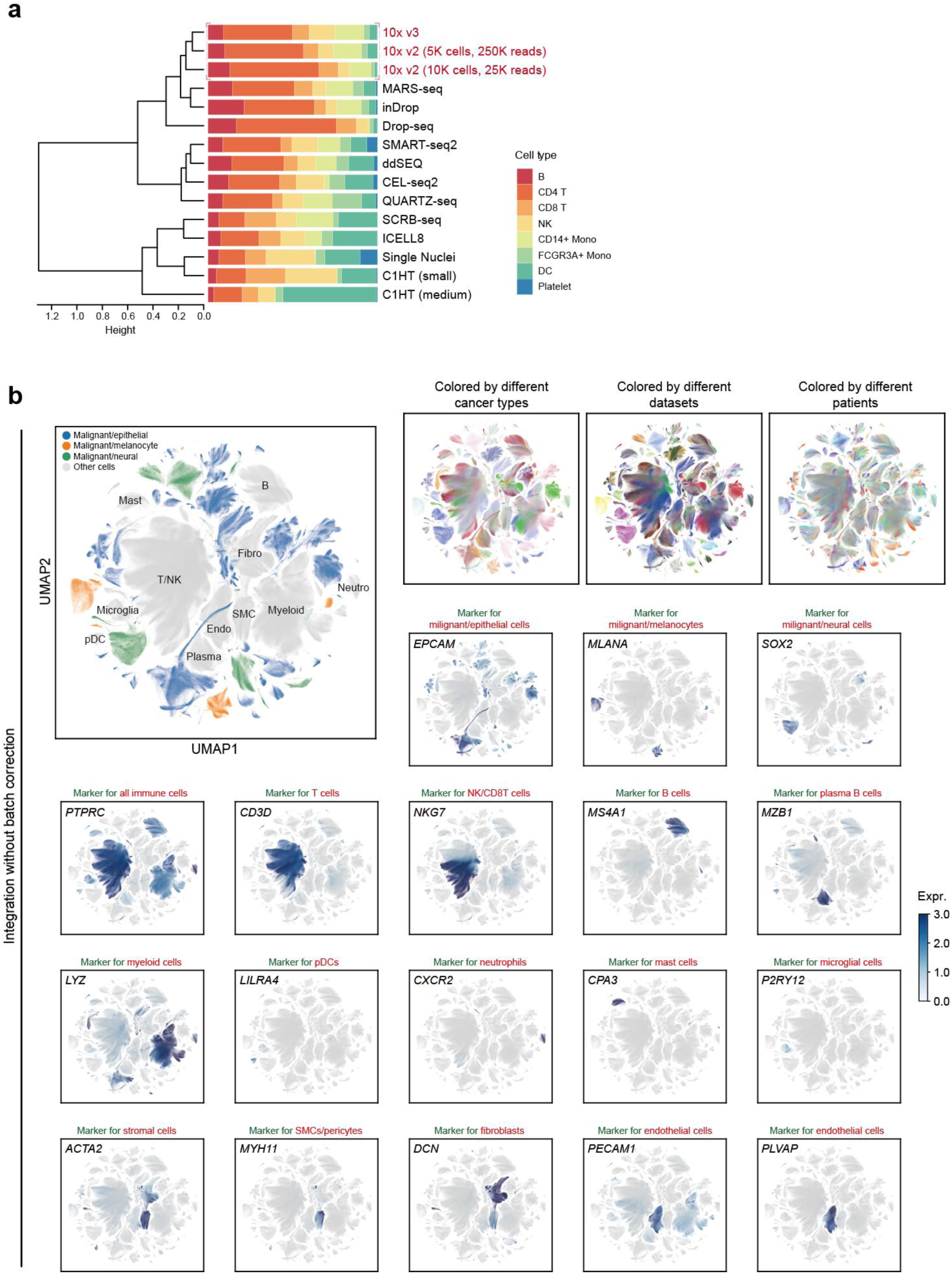
Evaluation of batch effects in scRNA-seq data. **a,** A comparison of the percentage of cell types detected in different single-cell RNA sequencing (scRNA-seq) platforms. Hierarchical clustering based on Euclidean distance is applied. **b,** UMAP embeddings of cell types without explicit batch correction during data integration, colored by different cancer types (left), datasets (middle), and patients (right). The expression of canonical markers of major cell types is displayed within the global (bottom) UMAP representation. The color gradient represents gene expression.

**Fig. S2.**
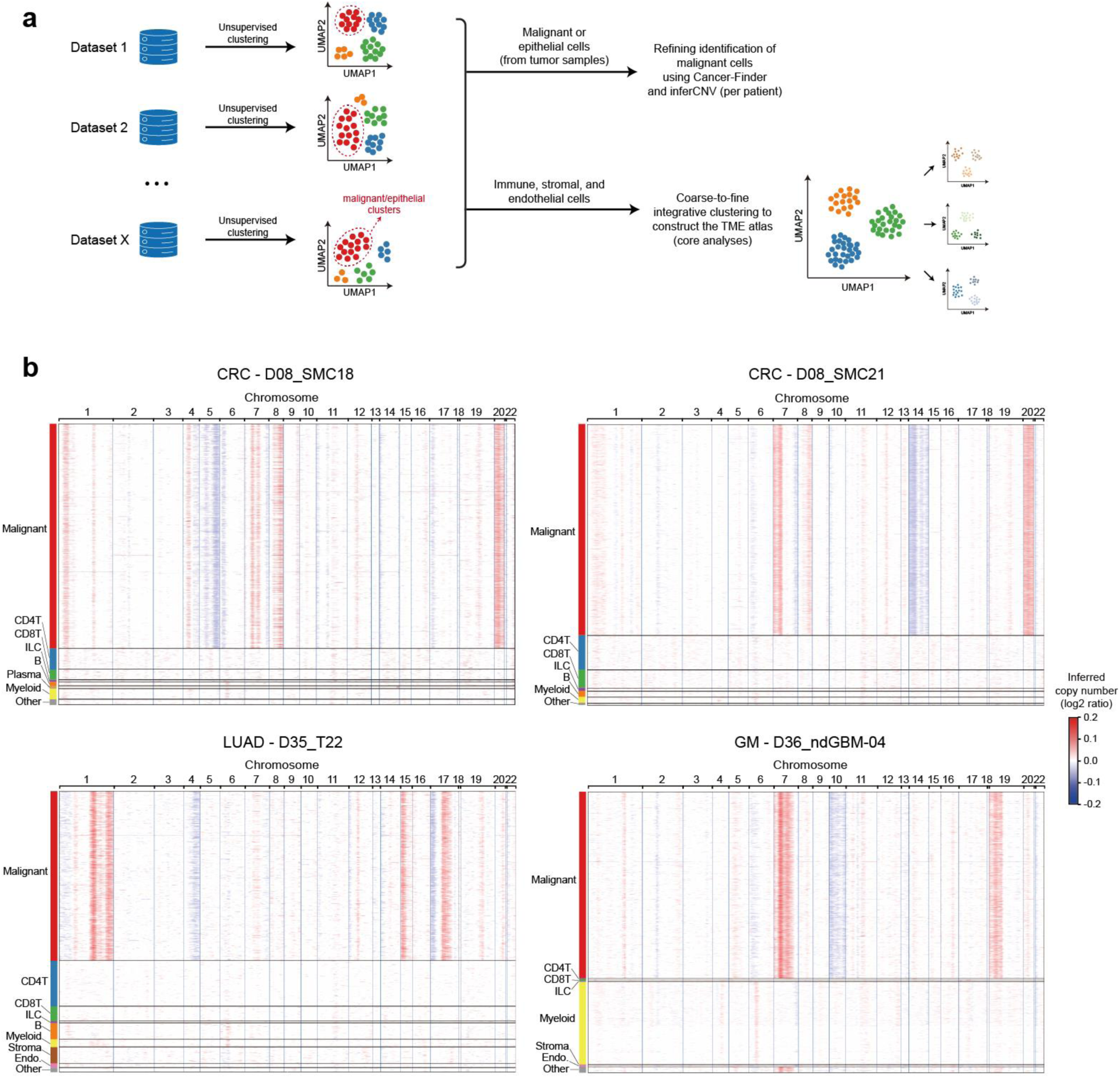
Analysis pipeline of scRNA-seq data and the identification of malignant cells. **a,** Schematic overview of the analysis pipeline of scRNA-seq datasets. **b,** Heatmap showing the representative CNV patterns in CRC (upper panel), LUAD (lower left), and GM (lower right) patients. Chromosomal regions are annotated at the top (for each column), while cell cluster identities are indicated by color bars on the left (for each row). CNV states are color-coded (loss: blue; gain: red).

**Fig. S3.**
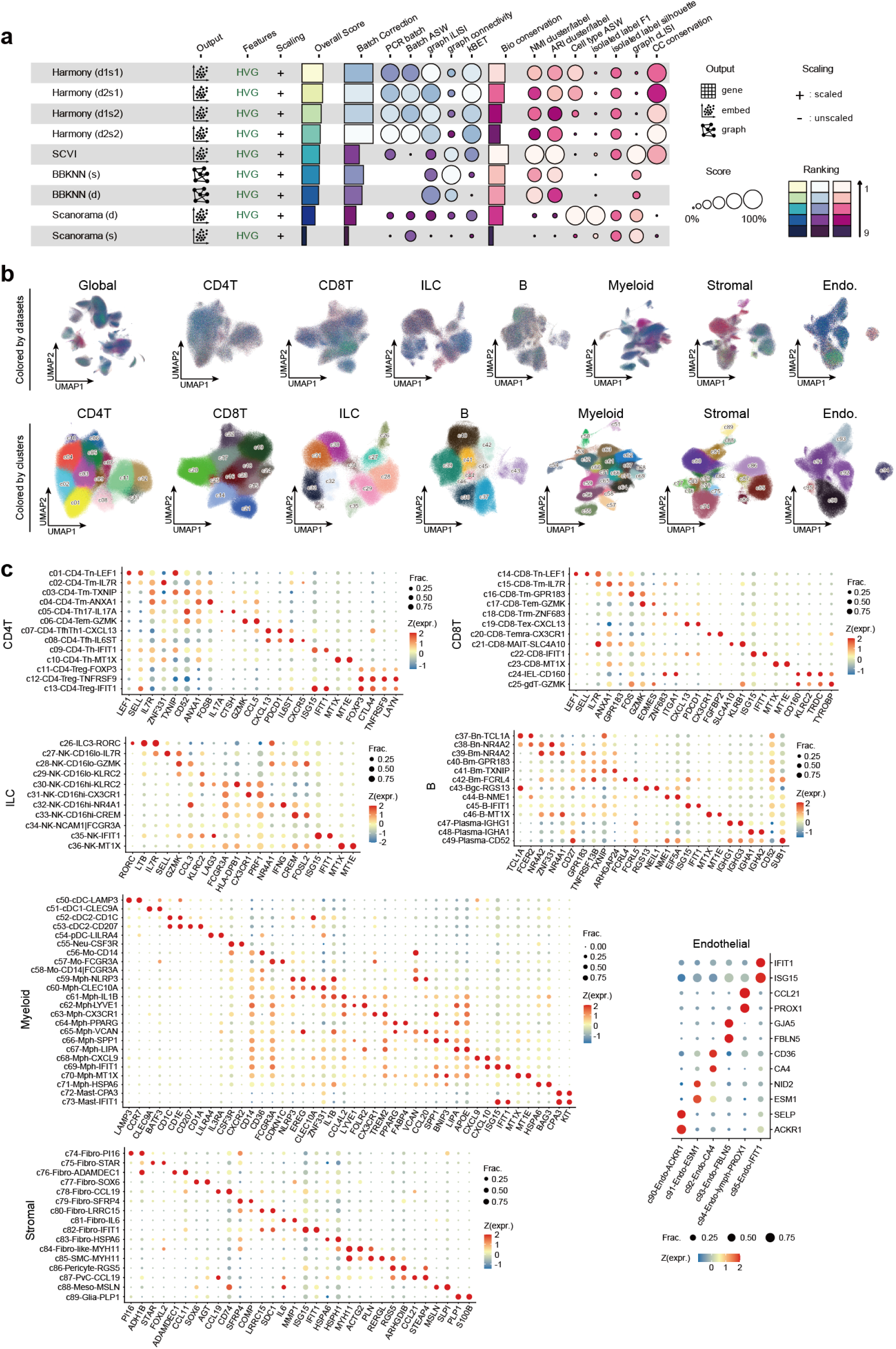
Integration analysis of scRNA-seq data. **a,** Performance of scRNA-seq integration methods was assessed by the single-cell integration benchmarking (scIB). Metrics include batch correction (blue) and bio-conservation (pink) categories. Overall scores are calculated by using a 40/60 weighted mean of these category scores. d, dataset; s, sample; 1/2, the value of theta. **b,** Visualizations of dataset information (top) and fine-grained sub-clusters (bottom)in the UMAP representation after Harmony integration. **c,** Dot plots showing the expression of representative signature genes of fine-grained sub-clusters of CD4^+^ T cells, CD8^+^ T cells, ILCs, B cells, myeloid cells, stromal cells, and endothelial cells. The dot size represents the proportion of expressing cells. The color indicates the z-score-scaled gene expression.

**Fig. S4.**
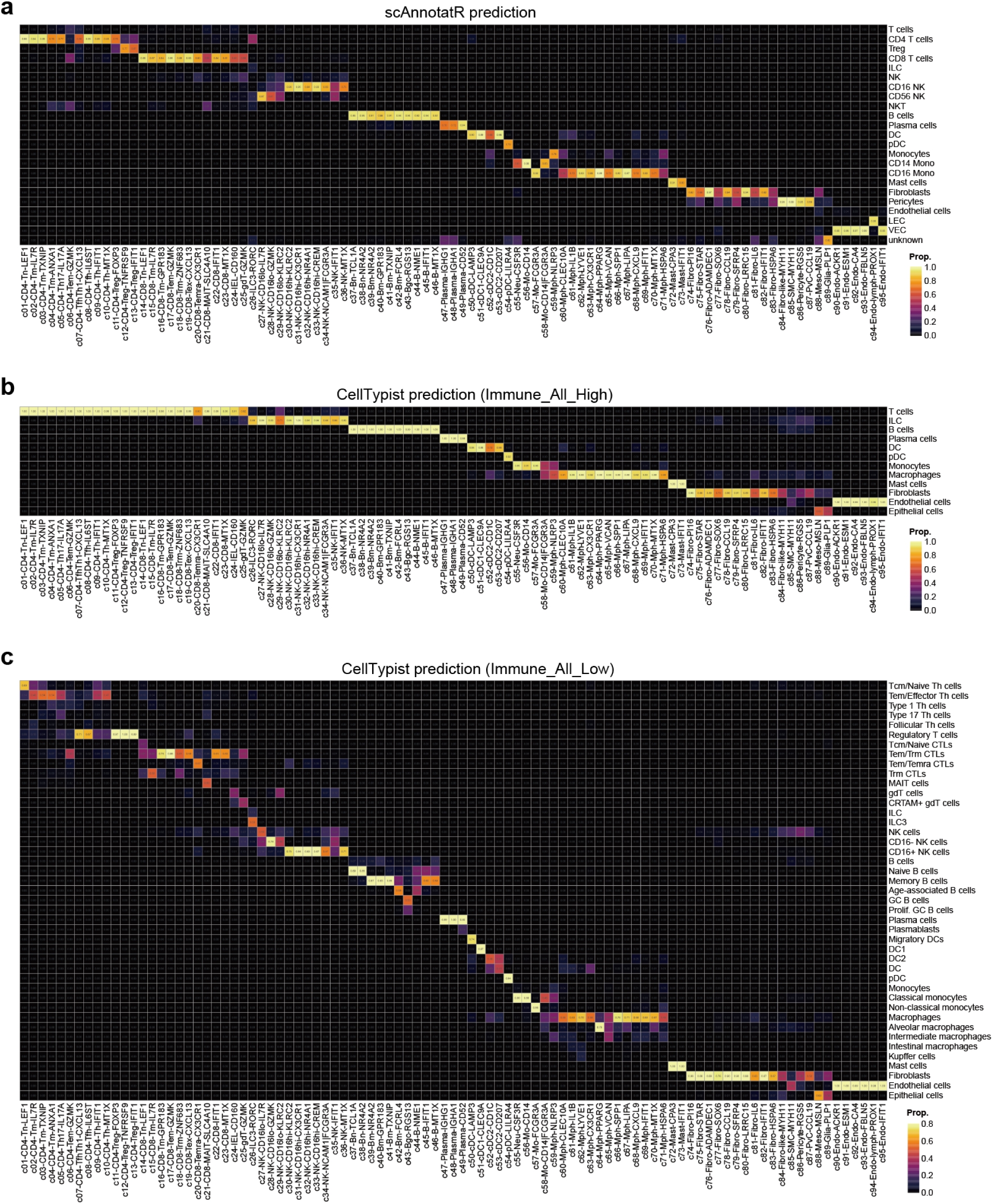
Validating the reliability of cell annotations. **a-c,** Heatmap comparing our cell annotations with predictions of scAnnotateR (**a**), CellTypist-Immune_All_High (**b**), and CellTypist-Immune_All_Low (**c**). Rows represent predicted cell types, columns denote annotated cell types, with color gradients indicating the proportion of each predicted cell type found in the corresponding annotated cell type. Only rows where the maximum proportion across all annotated cell types exceeds 0.05 are included.

**Fig. S5.**
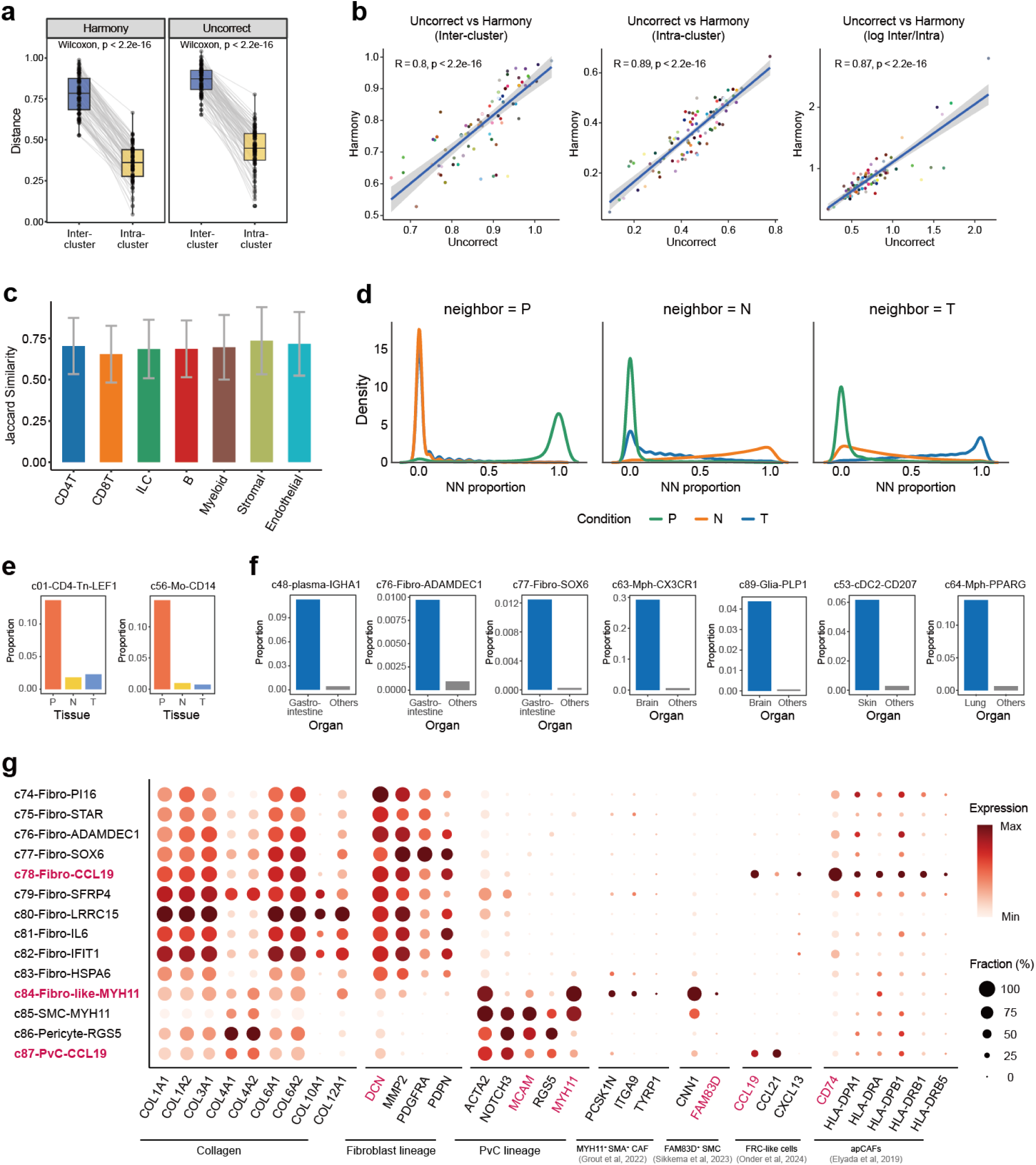
Integration quality assessment and features of rare stromal populations. **a,** Box plots showing the cosine distances of inter- and intra-cluster in both Harmony-corrected (left) and uncorrected data (right). Blue and yellow represent inter- and intra-cluster, respectively. Each dot represents a fine-grained sub-cluster. *P* values from paired Wilcoxon tests are shown. **b,** Scatter plots showing the Pearson correlation of inter-cluster distance (left), intra-cluster distance (middle), and the ratio of inter-cluster to intra-cluster distance in log transformation (right) between the uncorrected and Harmony-corrected data. Dots with different colors represent fine-grained sub-clusters. *R*, Pearson correlation coefficient. **c,** Bar plots depicting the Jaccard similarity of the 15 nearest neighbors within-batch for each cell between uncorrected and Harmony-corrected data. Colors indicate major clusters. Bar height represents the mean Jaccard similarity. Error bars represent standard deviation. **d,** Density plots for the distribution of out-of-batch nearest neighbors (oobNN) proportion from blood (P, left), normal (N, middle), and tumor tissues (T, right) around cells. **e,** Bar plots illustrating the distribution of sub-clusters enriched in blood. **f,** Bar plots showing the proportion of organ-specific sub-clusters. **g,** Dot plot showing the expression of representative signature genes of rare stromal populations. The dot size represents the proportion of cells expressing the genes. The color indicates the average level of gene expression.

**Fig. S6.**
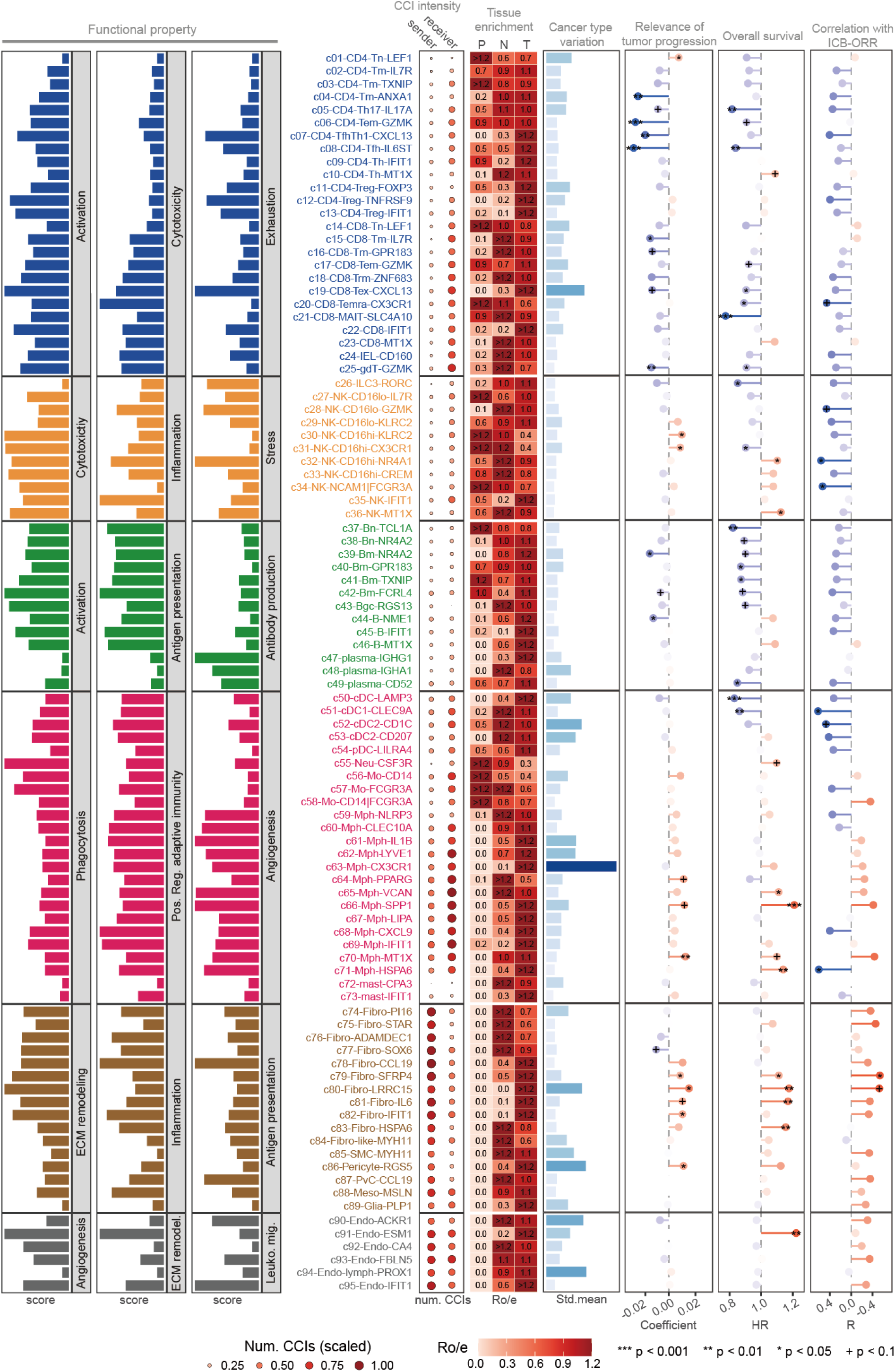
Summary of characteristics of fine-grained sub-clusters. This figure presents a summary of the characteristics of different sub-clusters. The first three columns of bar plots show the functional properties of each cell sub-cluster. Colors represent different major cellular compartments. The following dot plot displays the intensity of cell-cell interactions in each cell sub-cluster. Dot size and color denote scaled numbers of cell-cell interactions. The heatmap in the middle highlights the enrichment of cell sub-clusters in different tissues, which is calculated by the ratio of observed over expected cell numbers (Ro/e). The following bar plots separately demonstrate the variation in frequency and expression of cell sub-clusters across different cancer types. Lollipop charts for the last three columns show the association of cell sub-clusters with tumor progression, overall survival, and immune checkpoint blockade (ICB) objective response rates (ORRs), respectively. Blue represents clinically beneficial associations, and red represents clinically unfavorable associations. Significant associations are labelled with ‘+’ for *P* < 0.1, ‘*’ for *P* < 0.05, ‘**’ for *P* < 0.01 and ‘***’ for *P* < 0.001. Significance levels were computed via linear regression model, meta-analysis, and Pearson correlation test, respectively (details provided in the Methods section).

**Fig. S7.**
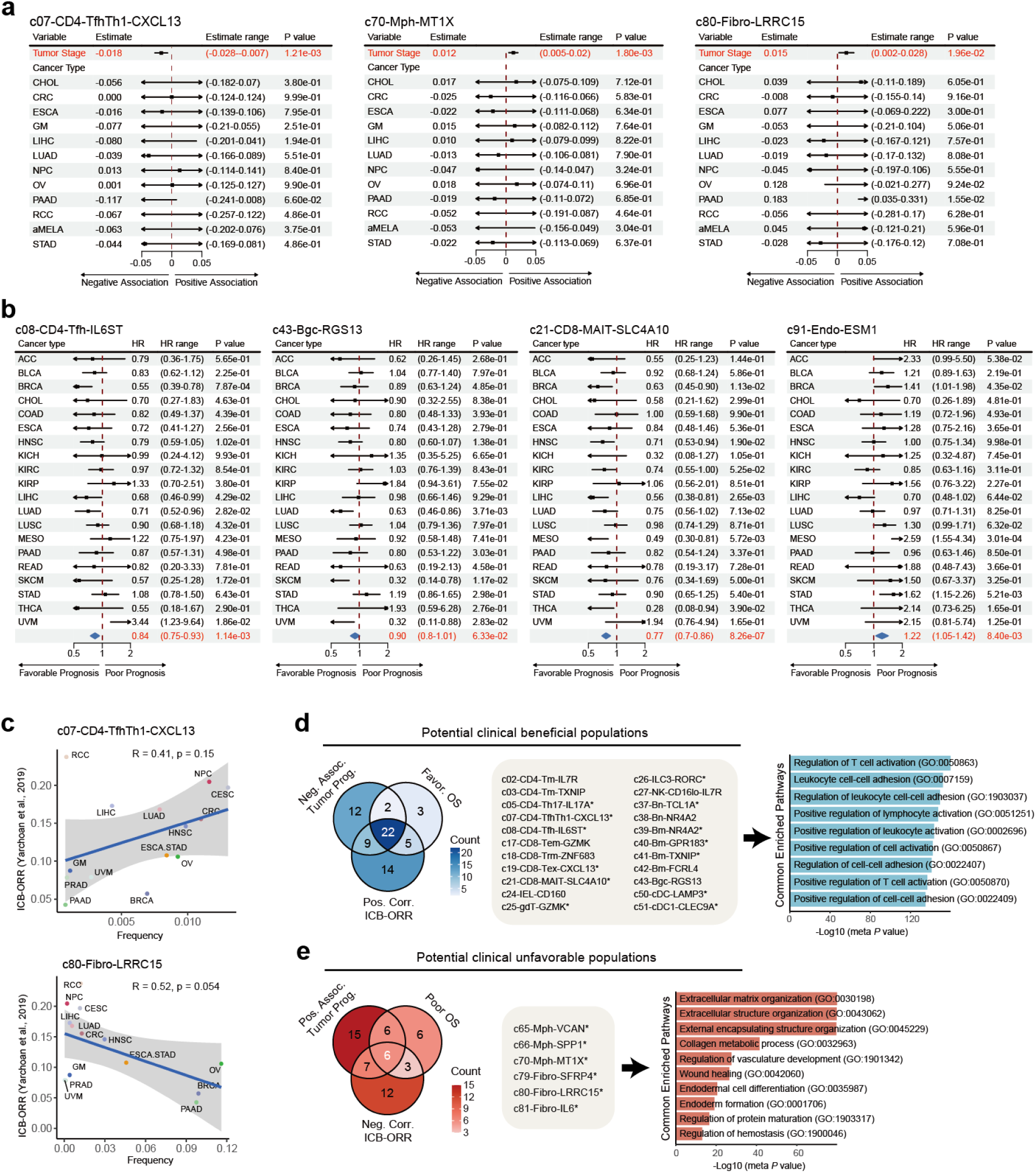
Examples of clinically relevant cell populations. **a,** Forest plots showing the relevance of tumor progression with the abundance of typical cell clusters. The estimated coefficients and their 95% confidence intervals, as well as the significance of the linear regression model, are shown in the forest plot. **b,** Forest plots showing the relationship between overall survival and the abundance of typical cell clusters. Arrows represent data that is out of range. HR, hazard ratio. Meta results are highlighted in red, and the hazard ratios are calculated using Cox regression models with the age, sex, and stage adjusted for in the forest plot. **c,** Scatter plots showing the immune checkpoint blockade objective response rates (ICB-ORRs) with the abundance of typical cell clusters. The Pearson correlation is measured and each dot represents a cancer type in the scatter plot. **d,** Venn diagram (left) illustrating the overlap among cell sub-clusters negatively associated with tumor progression, sub-clusters with favorable overall survival, and sub-clusters positively associated with ICB-ORRs. Color represents the number of cell sub-clusters. Sub-clusters achieving statistical significance in at least one clinically relevant analysis are marked with an asterisk (details provided in the Methods section). Bar plot (right) showing the common pathways of potential clinically beneficial populations (left) by gene ontology (GO) enrichment analysis. *P* values are calculated by the meta-analysis with the random effects model. **e,** Same as **(d)** for potentially clinically unfavorable populations.

**Fig. S8.**
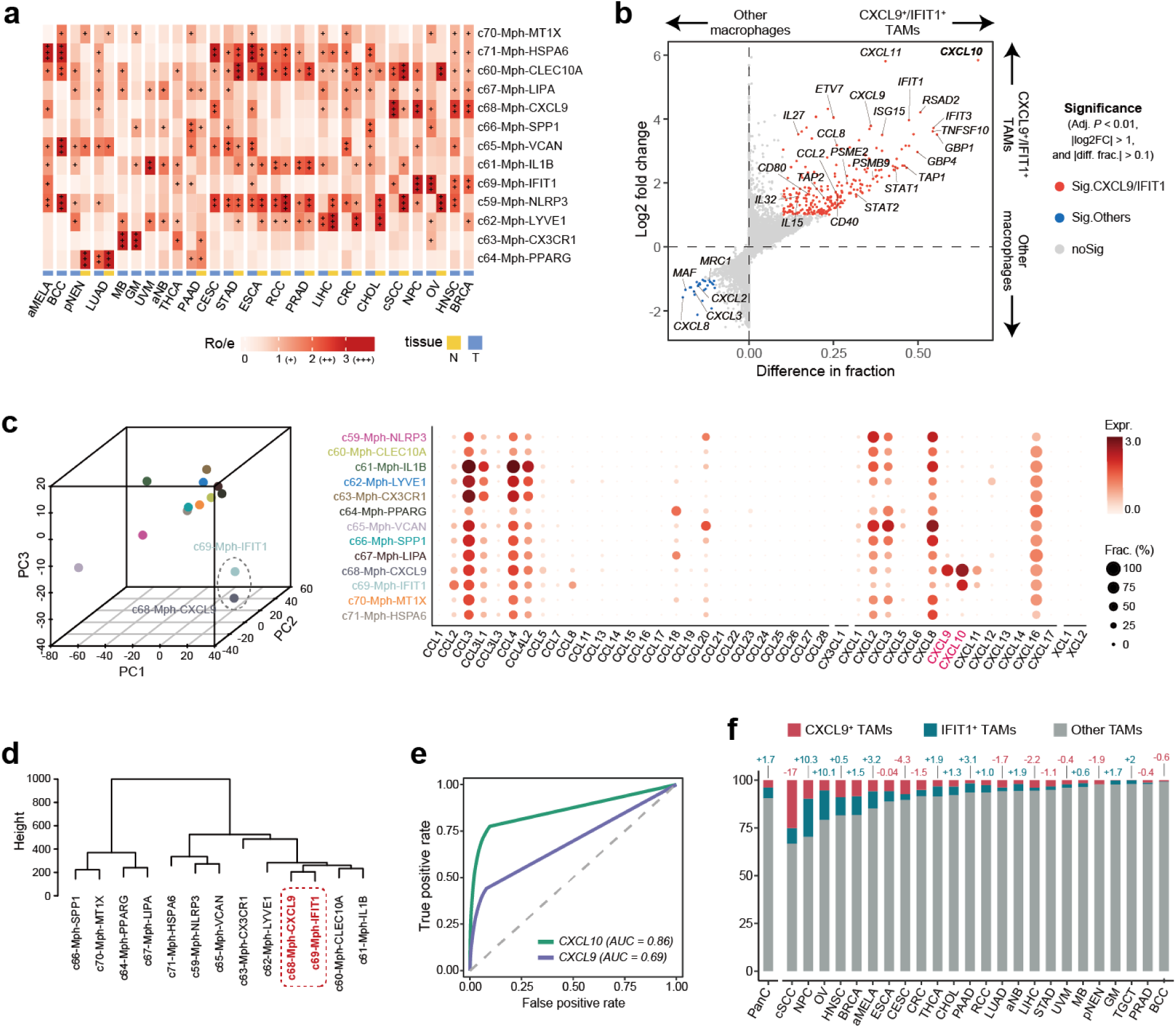
Heterogeneity of macrophage subsets. **a,** Heatmap showing the enrichment of cell sub-clusters in different cancer types, which is calculated by Ro/e. Yellow and blue represent non-tumor tissues (N) and tumor tissues (T), respectively. **b,** Scatter plot showing the differential expression of genes between *CXCL9*^+^/*IFIT1*^+^ and other TAMs. The x-axis represents the difference in detection fraction, while the y-axis displays log2FC. Genes with a BH-adjusted *P* value < 0.01 (two-sided Wilcoxon test), log2FC > 1, and difference in detection fraction > 0.1 are highlighted in red, while those with log2FC < -1 are highlighted in blue. **c,** 3D plots (left) of the principal component analysis (PCA) scores for the mean expression of chemokine-related genes in different macrophage clusters. Colors represent different clusters. Dot plot (right) showing the expression of chemokine-related genes in clusters of macrophages. The dot size represents the proportion of expressing cells. The color indicates the average level of gene expression. **d,** Hierarchical clustering of macrophage subsets. Euclidean distance was computed using the whole transcriptome profile. **e,** ROC plot showing the performance of selected genes in identifying *CXCL9*^+^ and *IFIT1*^+^ TAMs. Colors represent different selected genes. **f,** Bar plot showing the proportion of *CXCL9*^+^ and *IFIT1*^+^ TAMs across different cancer types. Red, green, and grey represent *CXCL9*^+^, *IFIT1*^+,^ and other TAMs, respectively. The numbers above the bars are differences between the proportion of *CXCL9*^+^ and *IFIT1*^+^ TAMs. Positive and negative numbers represent more *IFIT1*^+^ and more *CXCL9*^+^ TAMs, respectively.

**Fig. S9.**
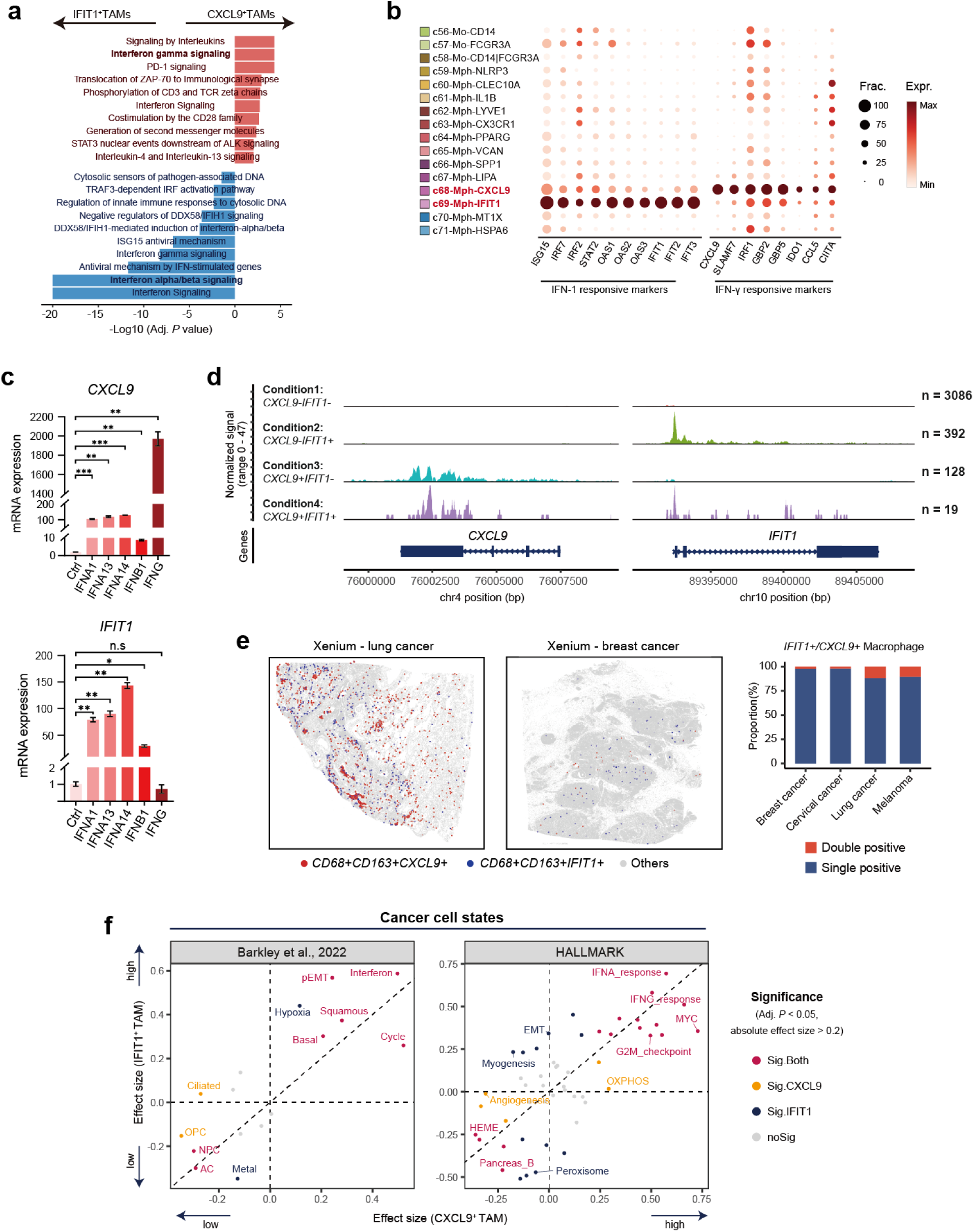
Distinction between *CXCL9*^+^ and *IFIT1*^+^ TAMs. **a,** GO enrichment analysis on genes differentially expressed in *CXCL9*^+^ and *IFIT1*^+^ TAMs. Red and blue represent pathways enriched in *CXCL9*^+^ and *IFIT1*^+^ TAMs, respectively. *P* values are adjusted by BH in the hypergeometric test. **b,** Dot plot showing the expression of representative signature genes of *CXCL9*^+^ and *IFIT1*^+^ TAMs. The dot size represents the proportion of expressing cells. The color indicates the average level of gene expression. **c,** Quantification of the mRNA expression of *CXCL9*^+^ macrophage-specific gene (*CXCL9*) and *IFIT1*^+^ macrophage-specific genes (*IFIT1*) in monocyte-derived macrophages, following a 24h stimulation with IFNA1, IFNA13, IFNA14, IFNB1, IFNG or control. *P* values are calculated using the two-sided t-test, where * indicates *P* < 0.05, ** indicates *P* < 0.01, and *** indicates *P* < 0.001. **d,** ATAC-seq tracks showing the chromatin accessibility in the *CXCL9* and *IFIT1* loci for macrophage subsets in TNBC tumors. **e,** Scatter plots showing the spatial distribution pattern of *CD68*^+^*CD163*^+^*CXCL9*^+^ cells and *CD68*^+^*CD163*^+^*IFIT1*^+^ cells in representative Xenium samples. Each dot represents an individual cell, and color represents the cell identity. The right bar plots showing the proportion of *IFIT1*^+^ or *CXCL9*^+^ macrophages across Xenium samples. **f,** Scatter plots showing effect sizes of the cancer cell state comparisons between c68-CXCL9^high^ versus c68-CXCL9^low^ and c69-IFIT1^high^ versus c69-IFIT1^low^ tumors. Left: recurrent cancer gene programs (Barkley et al.); right: HALLMARK pathways (MsigDB). Dots represent cancer cell states, and colors represent statistical significance categories. Effect sizes are calculated as Hedge’s g values derived from Student’s t-test, and *P* values are adjusted by the BH-method. Dots with a BH-adjusted *P* value < 0.05 and an absolute effect size >0.2 are highlighted.

**Fig. S10.**
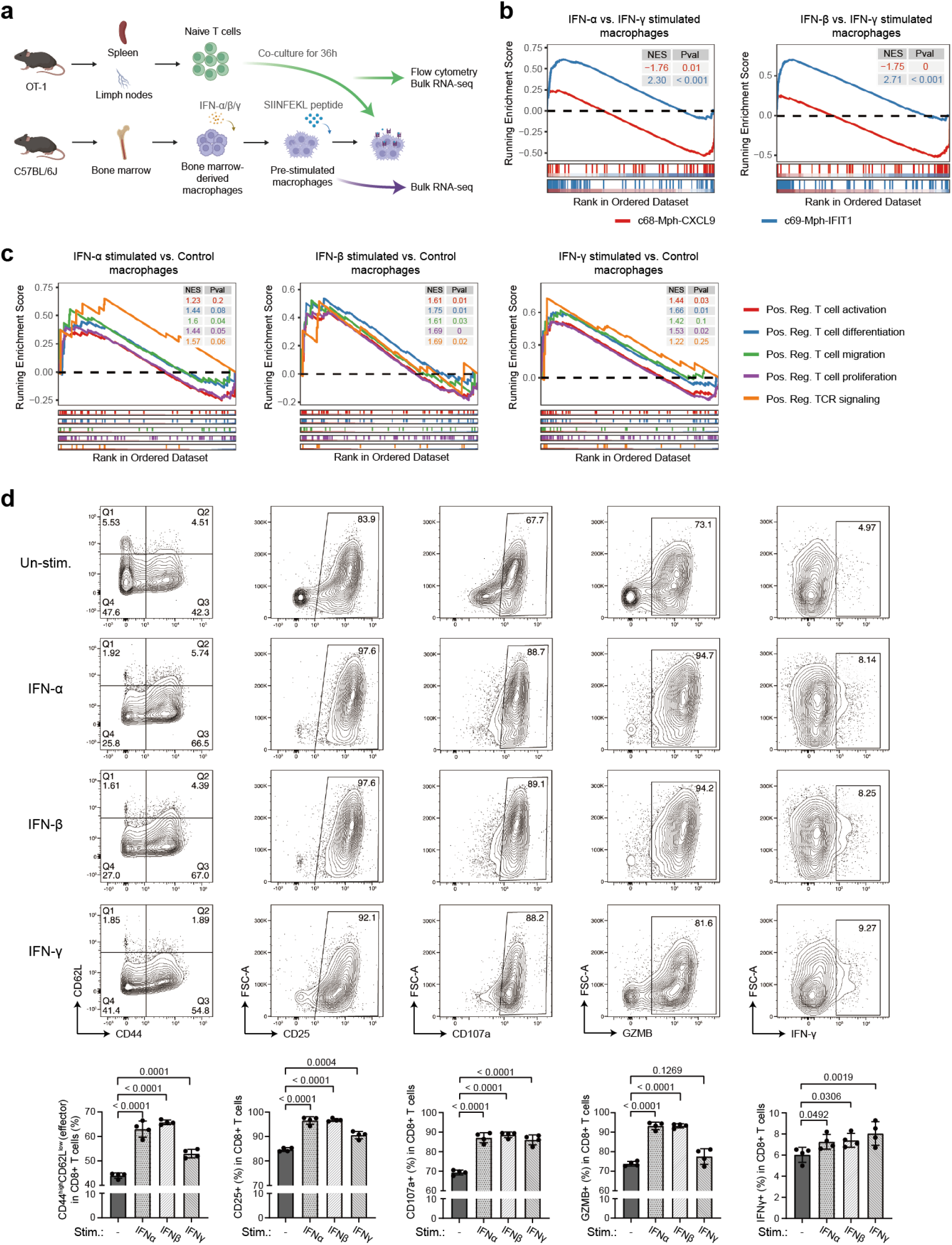
*In vitro* experiments for validating the functions of *CXCL9*^+^ and *IFIT1*^+^ TAMs. **a,** Schematic diagram of the co-culture experiments. **b,** Gene Set Enrichment Analysis (GSEA) comparing IFN-α stimulated versus IFN-γ stimulated macrophages (left), and IFN-β stimulated versus IFN-γ stimulated macrophages (right). Colors represent gene sets (top 100 signature genes) from c68-Mph-CXCL9 (red) and c69-Mph-IFIT1 (blue). **c,** GSEA comparing IFN-stimulated versus unstimulated macrophages: IFN-α (left), IFN-β (middle), IFN-γ (right). Colors correspond to different gene sets from MSigDB. **d,** Representative flow cytometry plots and percentages of CD44^+^CD62^-^/ CD25^+^/ CD107a^+^/GZMB^+^/IFN-γ^+^ CD8 T cells following 36-hour co-culture with IFN-stimulated BMDMs (n=4). The *P* value is calculated by the two-sided t-test.

**Fig. S11.**
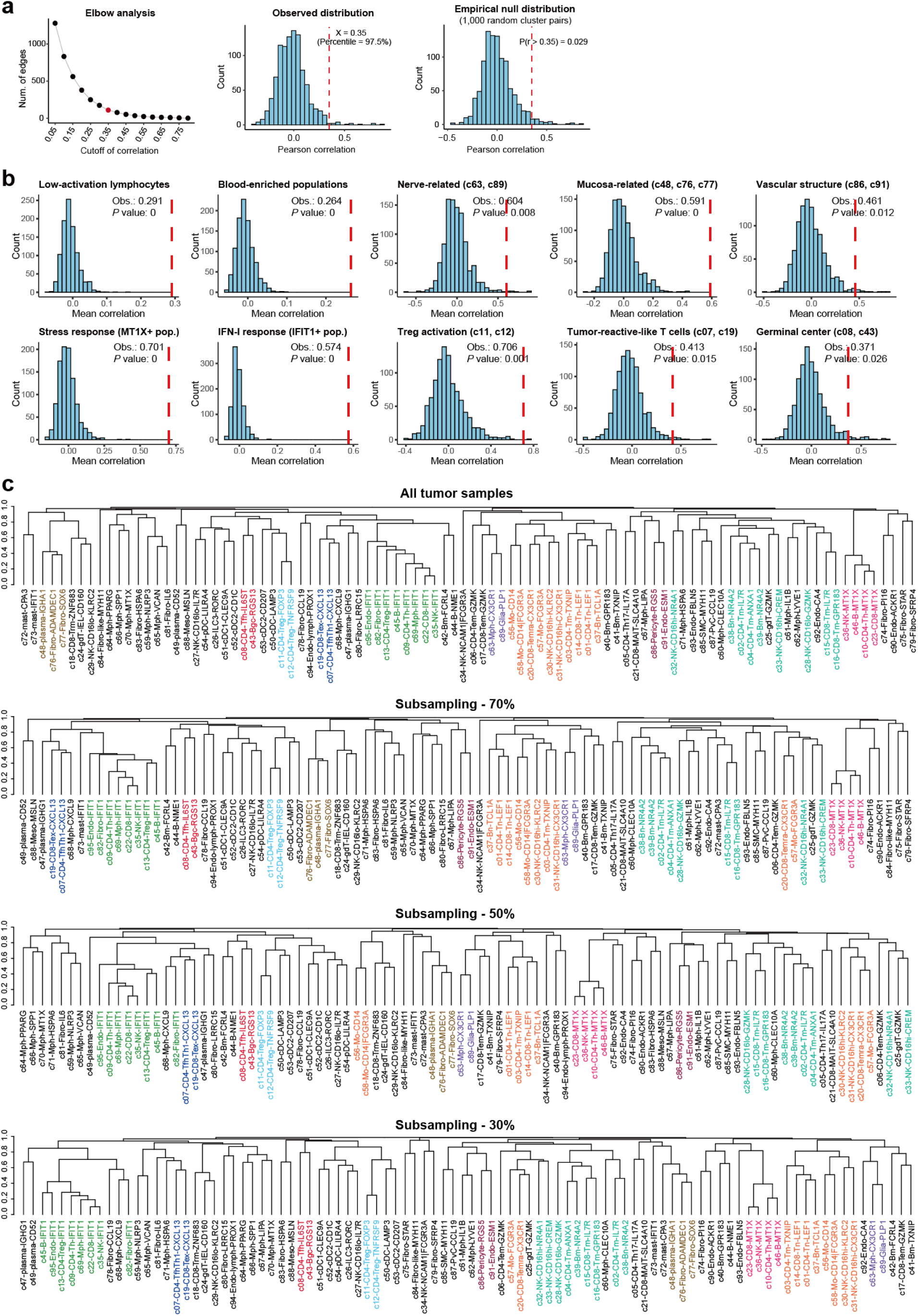
Identification and robustness analysis of co-occurrent patterns. **a,** Evaluating the correlation cutoff for constructing the co-occurrence network. The first (left) line graph showing the number of edges under different thresholds. The knee point is marked in red. The following (middle) histogram plot showing the distribution of all pairwise correlation coefficients between cell clusters across the pan-cancer data. The cutoff of 0.35 is marked by the red dashed line. The last (right) histogram plot showing the empirical null distribution from 1,000 randomized cell cluster pairs, and the cutoff of 0.35 is marked by the red dashed line. **b,** Histogram plots showing the average correlation among cell clusters within modules and the distribution of 1,000 random permutation tests. The red dotted line represents the observed intra-module correlation. *P* values are calculated as the proportion of random correlations exceeding the observed intra-module correlations. **c,** Dendrogram plots showing the hierarchical clustering of core datasets with 100%, 70%, 50%, and 30% of the data retained. Cell clusters with the same color belong to the same module.

**Fig. S12.**
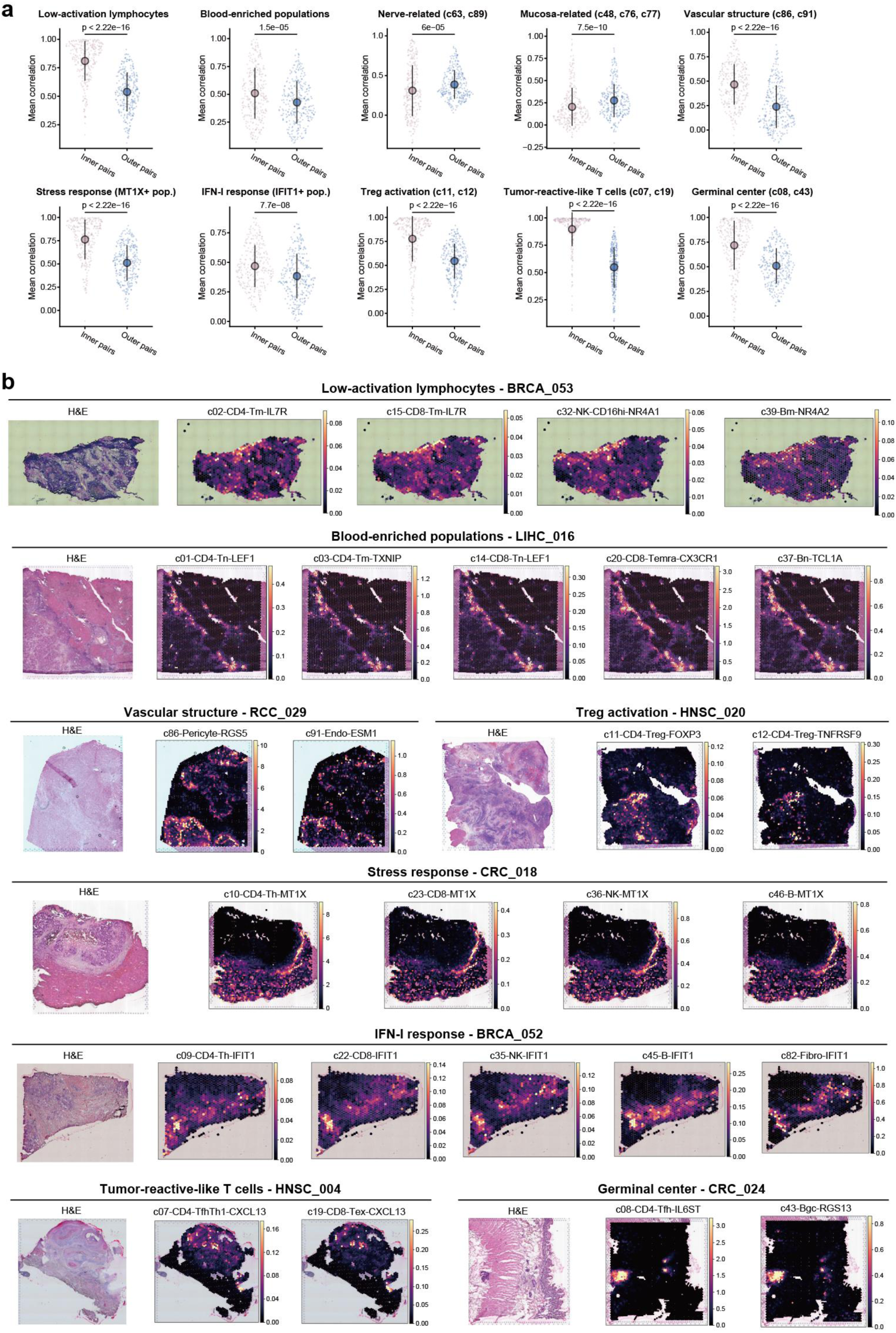
Quantitative assessment and visual representation of the spatial distribution of co-occurrent cluster pairs. **a,** Sina plots comparing the mean Pearson correlations of cell clusters within the modules (inner pairs) and outside the module (outer pairs). Each dot represents one sample. The *P* value is calculated by the two-sided Wilcoxon test. **b,** Representative spatial plots showing the spatial distribution of cell clusters within a specific co-occurrent module. Color represents the relative abundances predicted by cell2location.

**Fig. S13.**
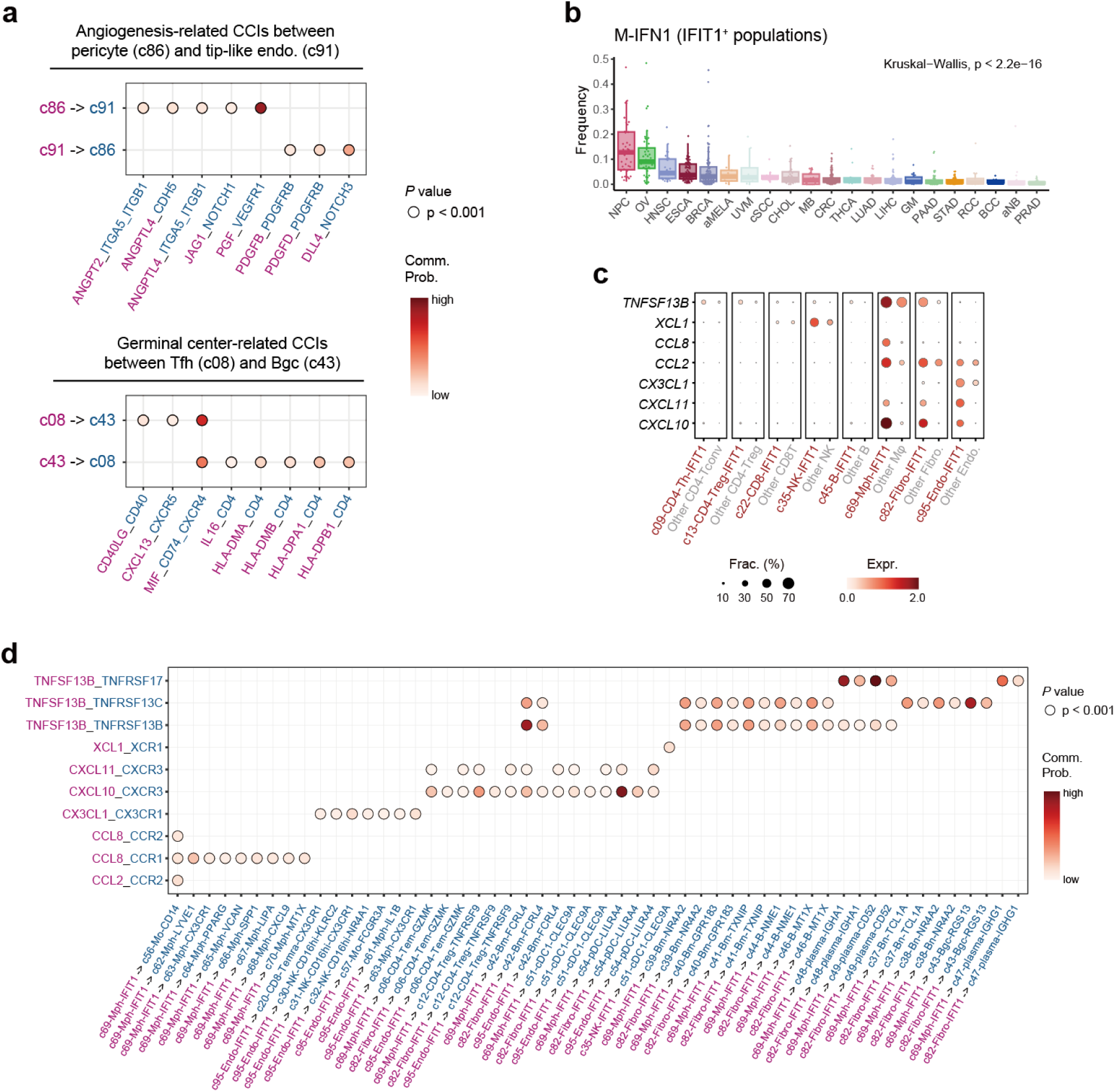
Characteristics of co-occurrent cluster pairs. **a,** Bubble heatmap showing the cell-cell communication of angiogenesis (left) and germinal center (right) related ligand-receptor pairs between specific sub-clusters. Dot sizes indicate *P* values, and color represents the communication probability. **b,** Box plot showing the M-IFN1 (*IFIT1*^+^ populations) distribution among cancer types. *P* value is calculated using a Kruskal-Wallis test. **c,** Dot plot showing the upregulated expression of chemokine and cytokine ligands in M-IFN1 clusters. The dot size represents the proportion of expressing cells. The color indicates the gene expression level. **d,** Bubble heatmap showing the cell-cell communication of specific ligand-receptor pairs (Fig. 3d) between sub-clusters of *IFIT1*^+^ populations and other TME cells. Dot sizes indicate *P* values, and color represents the communication probability.

**Fig. S14.**
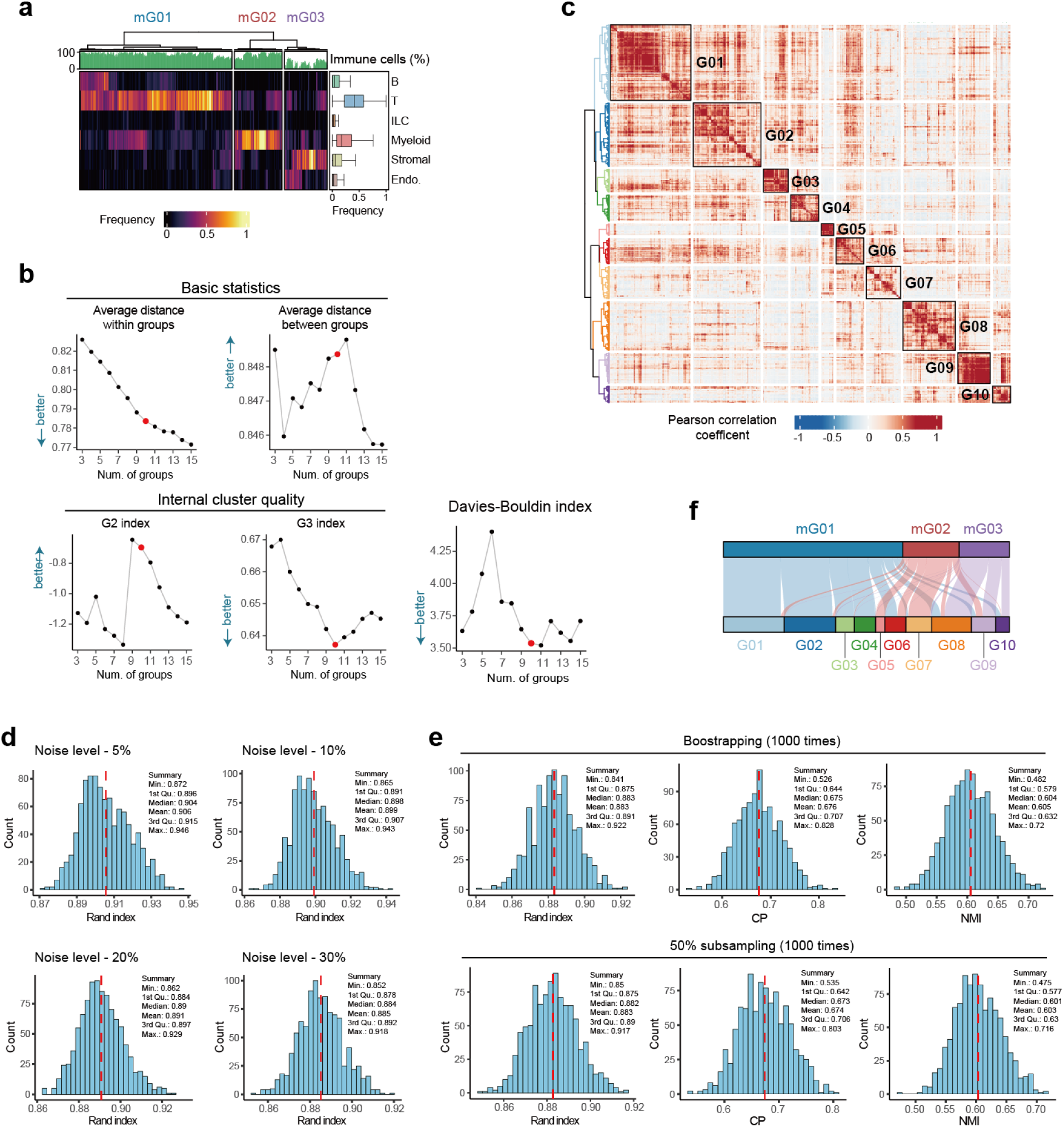
Identification and robustness analysis of 10 fine-grained TME groups. **a,** Heatmap illustrating frequencies of six major cell compartments in tumors. The rows correspond to the major cell compartments, and the columns represent tumor samples. For rows, box plots illustrate the distribution of major cell compartments across samples; for columns, bar plots illustrate the proportion of immune cells. Hierarchical clustering based on Pearson distance is applied to columns. **b,** Connected scatter plots showing changes in quality metrics when different numbers of groups are defined. The point corresponding to 10 groups is highlighted in red. **c,** Heatmap showing the Pearson correlation of patients based on the frequencies of fine-grained sub-clusters in tumors. Color represents the Pearson correlation coefficient. **d,** Histogram plots showing the distribution of Rand index between the original TME groups and the perturbed TME groups, which were generated by adding stochastic Gaussian noise at 5%, 10%, 20% and 30%. The red dotted line represents the mean Rand index. **e,** Histogram plots showing the distribution of Rand index, clustering purity (CP), and normalized mutual information (NMI) between the original TME groups and the perturbed TME groups generated by bootstrapping (top) and 50% subsampling (bottom). The red dotted lines represent the mean values. **f,** Sankey plot showing the relationship between tumor classifications by major compartments and by sub-clusters.

**Fig. S15.**
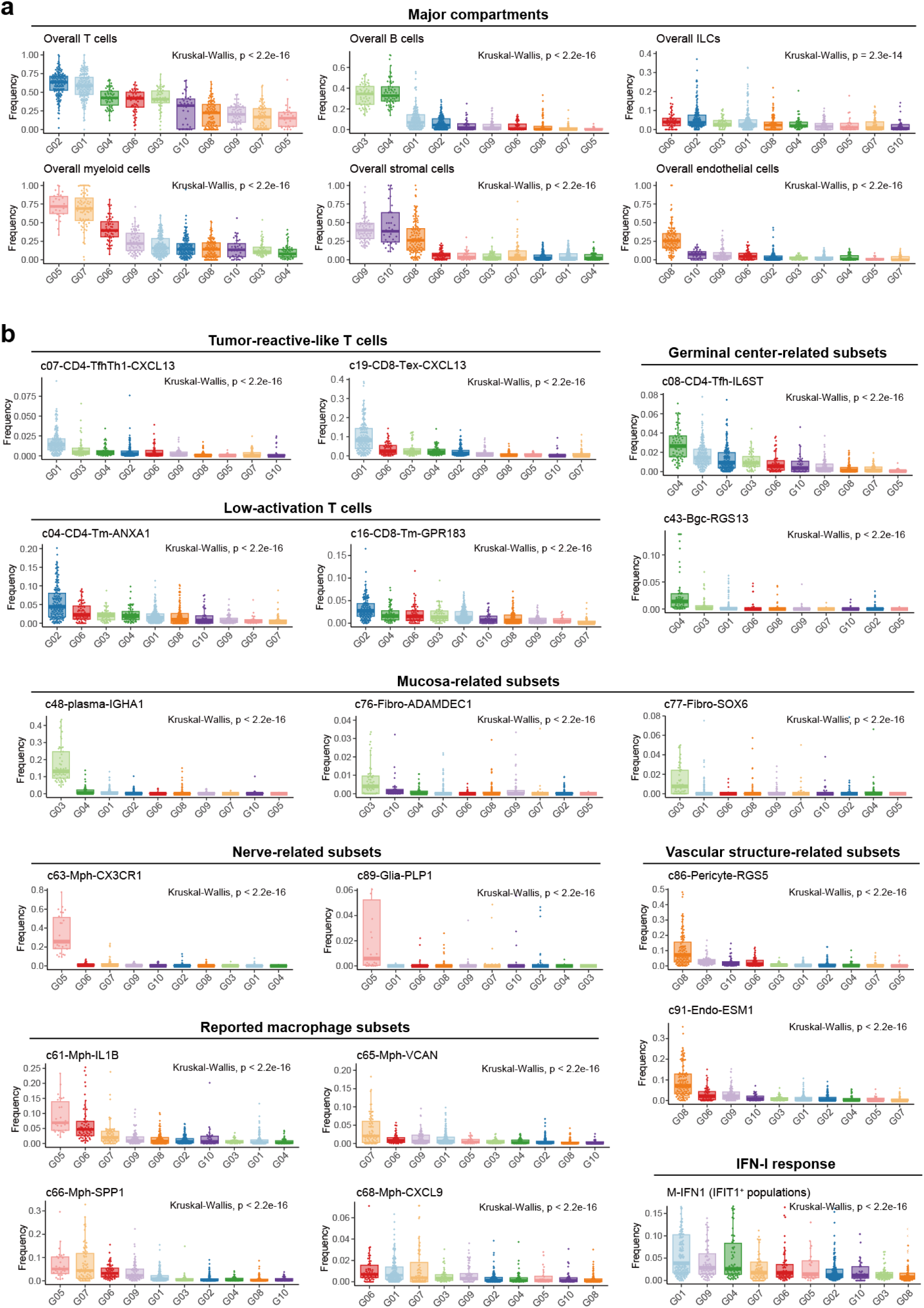
The distribution of cell clusters in TME groups. **a-b,** Box plots comparing frequencies of six major compartments **(a)** and representative cell sub-clusters **(b)** among different TME groups. *P* values calculated by the Kruskal-Wallis test are shown.

**Fig. S16.**
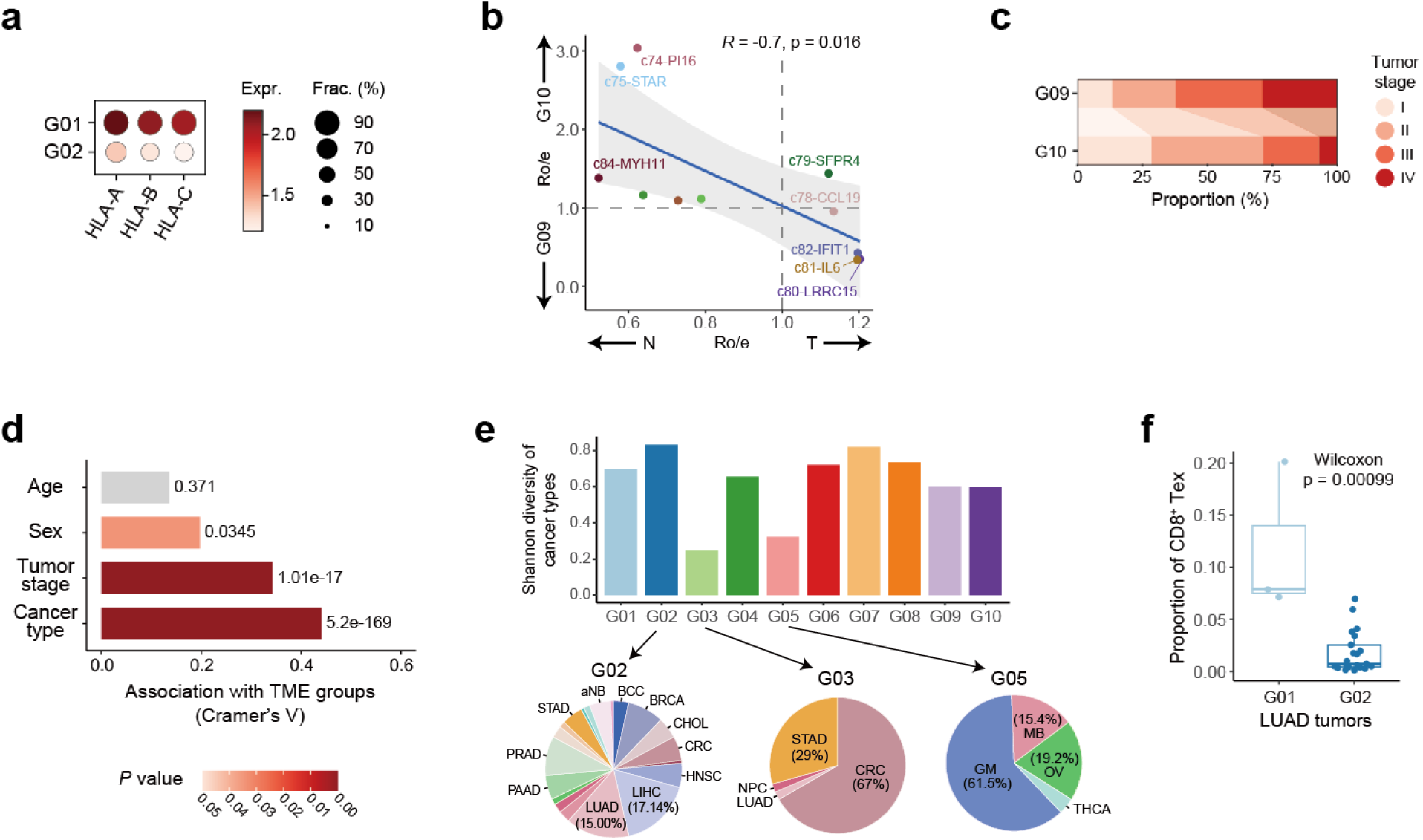
Characteristics of TME groups. **a,** Dot plot showing the expression of *HLA-A/B/C* in cancer cells derived from G01 and G02 tumors. The dot size represents the proportion of cells expressing genes. The color indicates the mean gene expression level. **b,** Scatter plot showing the Pearson correlation of Ro/e values between TME groups (G09 and G10) and tissues (N and T). Dots with different colors represent fibroblast sub-clusters. *R*, Pearson correlation coefficient. **c,** Bar plots showing the proportion of tumors with different stages in G09 and G10. Colors represent different tumor stages. **d,** Bar plot showing the association between the clinical parameters of patients and TME groups. The color indicates the *P* value calculated by the chi-square test. **e,** Bar plot showing the Shannon diversity of cancer types within each TME group. Colors represent different TME groups. Pie plots showing the examples of cancer type proportion in G02, G03, and G05. **f,** Box plot comparing the Tex cell frequency between LUAD tumors grouped in G01 and those in G02. The *P* value is calculated by the two-sided Wilcoxon test.

**Fig. S17.**
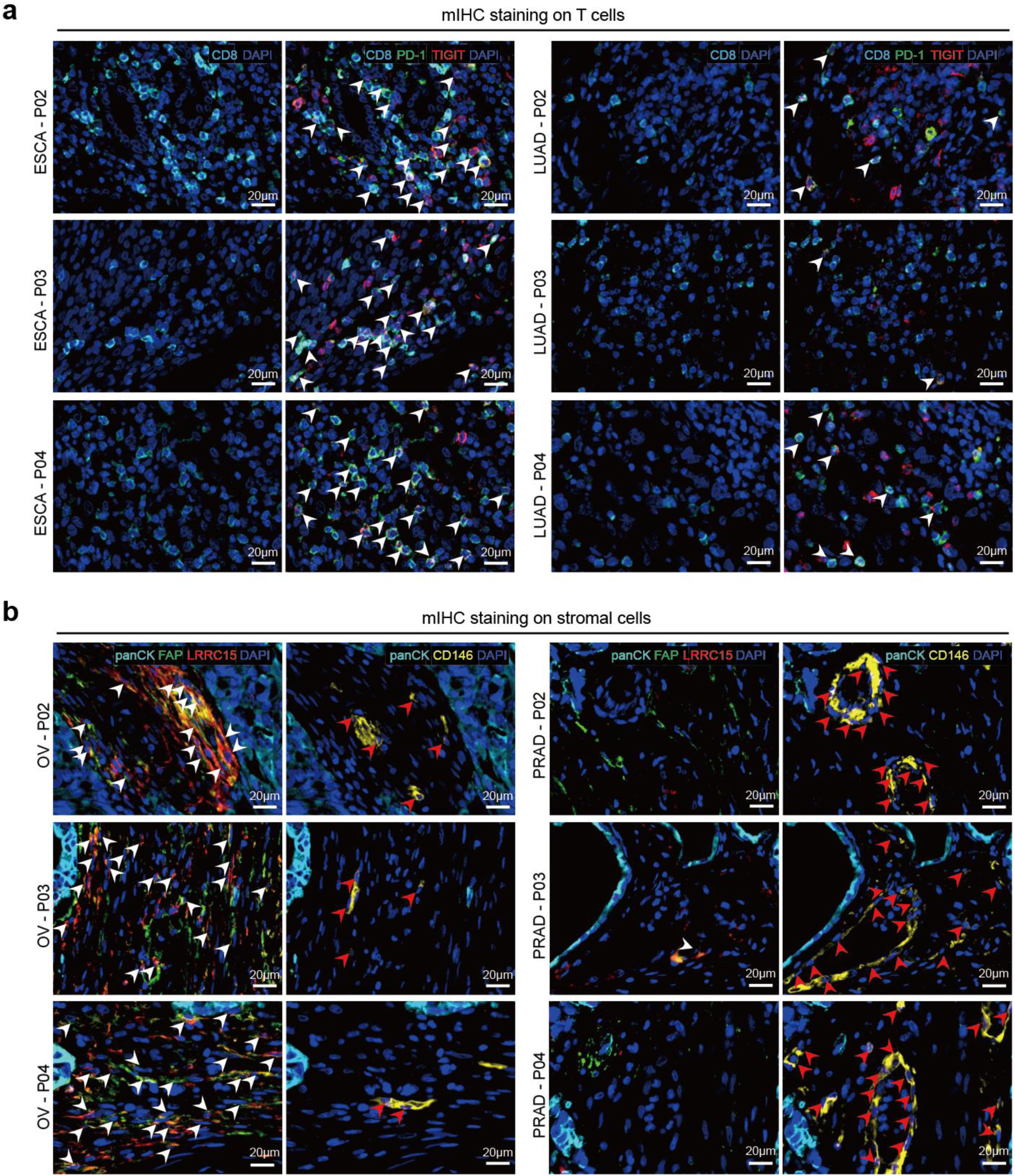
Representative views of multiplex immunofluorescence staining. **a,** Multiplex immunofluorescence staining on CD8^+^ T cells in ESCA (left columns) and LUAD (right columns) patients with anti-CD8 (blue), PD1 (green), and TIGIT (red) antibodies. **b,** Multiplex immunofluorescence staining on Stromal cells in OV (left columns) and PRAD (right columns) patients by anti-FAP (green), LRRC15 (red), and CD146 (yellow) antibodies. Scale bar, 20 μm.

**Fig. S18.**
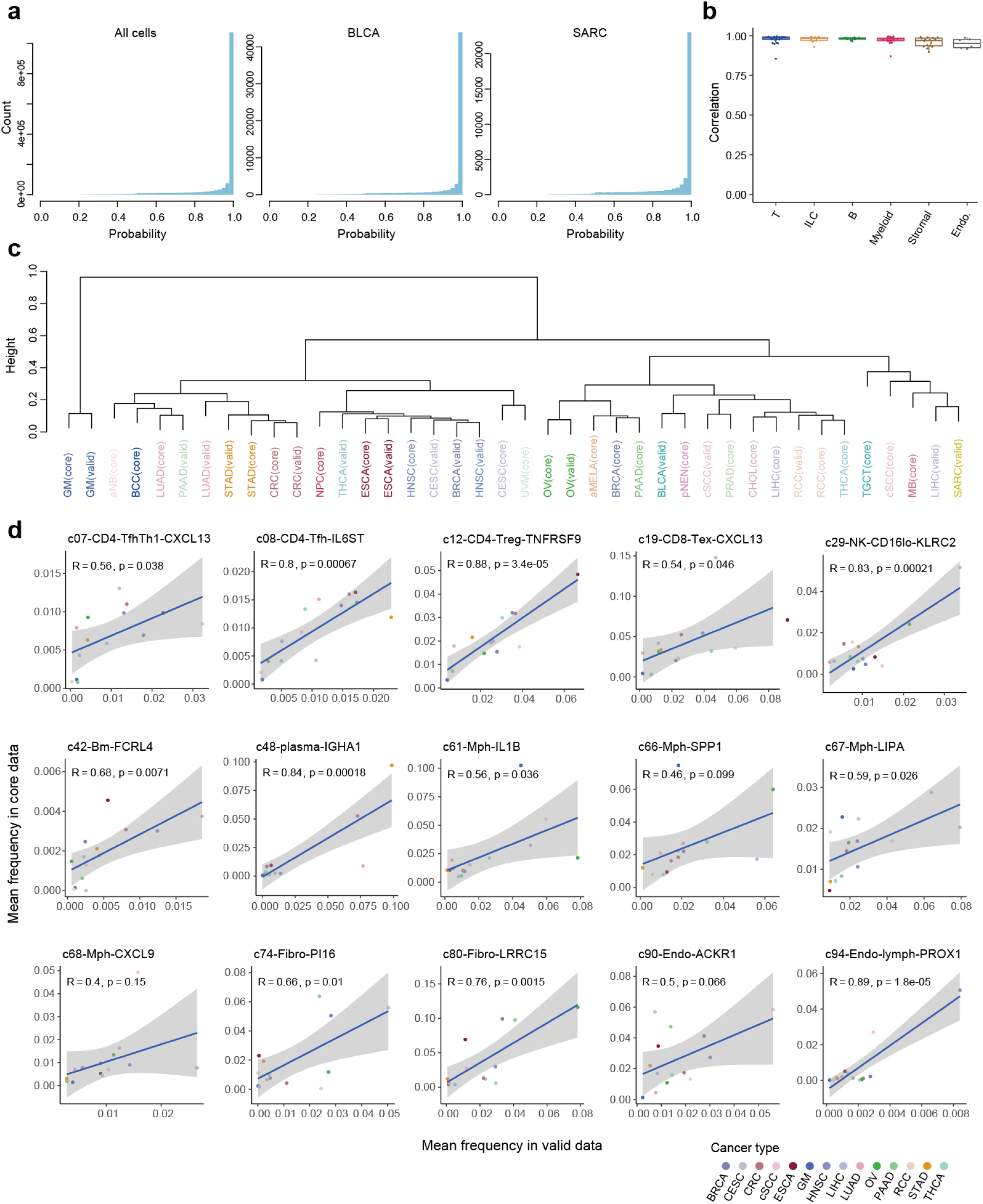
Analysis of the validation set at the cell-cluster level. **a,** Histogram showing the distribution of TOSICA prediction probability for cells from the entire validation set (left) and cells from BLCA and SARC patients (right). **b,** Box plot showing correlations between the transcriptome profiles of sub-clusters within the validation set and their counterparts in the core set. Colors represent different cell compartments. **c,** Hierarchical clustering of cancer types in different cohorts based on the Euclidean distance of their sub-cluster compositions. Colors represent different cancer types. **d,** Scatter plots showing the Pearson correlations between the mean frequencies of representative sub-clusters in each cancer type of the validation set and the corresponding values in the core set. Colors represent different cancer types. Only cancer types present in both cohorts are shown.

**Fig. S19.**
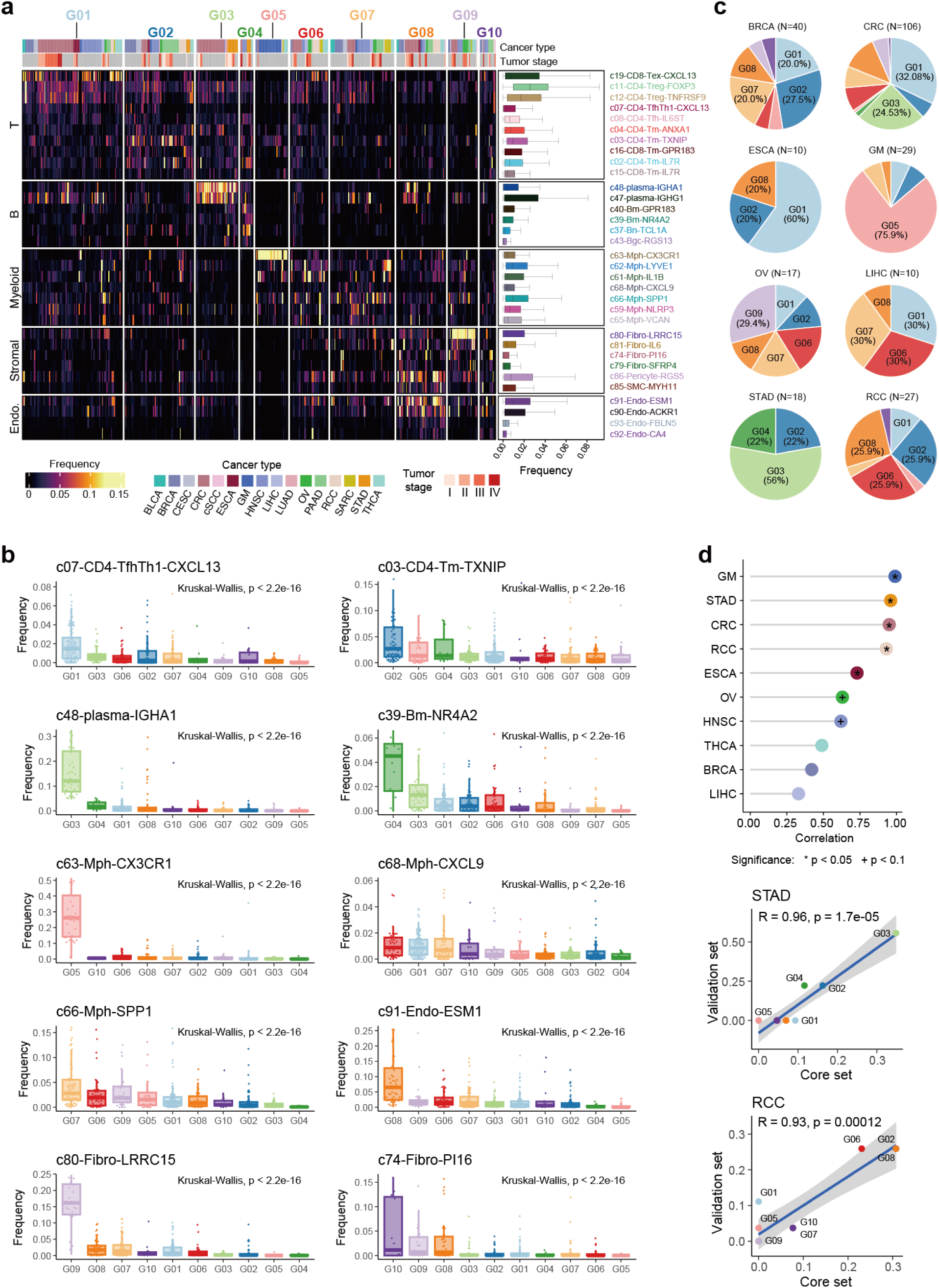
Analysis of the validation set at the TME-group level. **a,** Heatmap showing frequencies of predicted sub-clusters in tumors of the validation set. For rows, box plots illustrate the distribution of predicted sub-clusters across samples. **b,** Box plots comparing frequencies of representative sub-clusters among different TME groups in the validation set. *P* values calculated by the Kruskal-Wallis test are shown. **c,** Pie charts showing proportions of TME groups among different cancer types in the validation set. Colors represent TME groups. **d,** Lollipop plot (top) illustrating the Pearson correlation of TME group compositions between validation and core sets. Significant associations are labelled with ‘+’ for *P* < 0.1 and ‘*’ for *P* < 0.05. Only cancer types present in both sets and with >20 patients in the core sets are included. Scatter plots (bottom) showing the Pearson correlation of TME group compositions between validation and core sets in STAD and RCC.

**Fig. S20.**
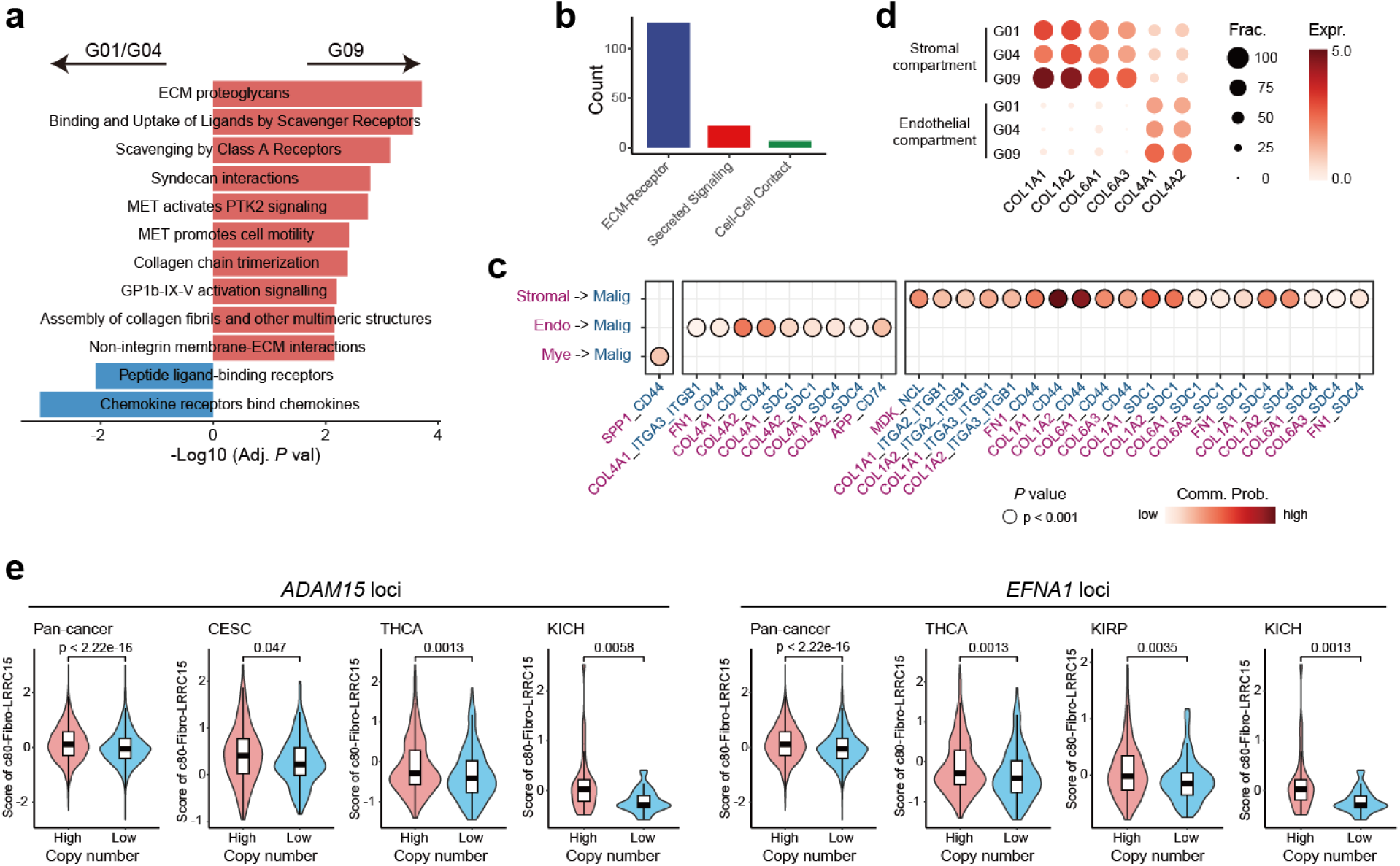
Functional and intercellular communication differences between G01/G04 and G09. **a,** GO enrichment analysis of genes differentially expressed in G09 and G01/G04 tumors. Red and blue represent pathways enriched in G09 and G01/G04 tumors, respectively. *P* values are adjusted using BH method in the hypergeometric test. **b,** Bar plot showing the count of different cell-cell interaction (CCI) types in G09. **c,** Dot plots showing the selected CCIs in G09. Colors represent the communication probability, and dot sizes reflect *P* values. **d,** Dot plot showing the expression of collagen IV and collagen I/VI genes in tumors from G01, G04, and G09. The dot size represents the proportion of expressing cells. The color indicates the gene expression level. **e,** Violin plots showing scores of c80-Fibro-LRRC15 across tumors grouped by the copy numbers of *ADAM15* (left) and *EFNA1* (right), at the pan-cancer level and in representative cancer types. The *P* value is calculated using the two-sided Wilcoxon test.

**Fig. S21.**
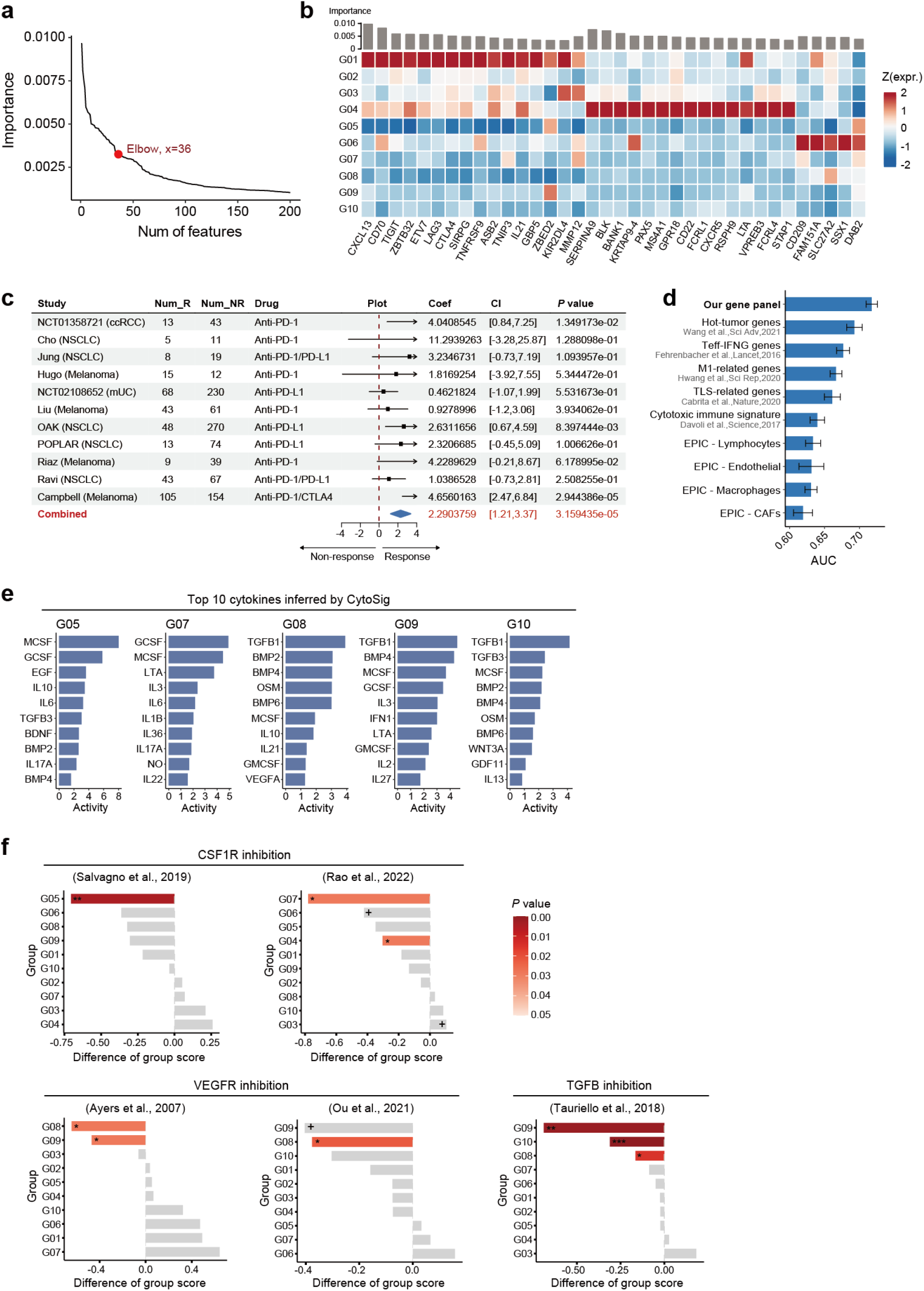
Therapeutic associations of stratified TME groups. **a,** Line chart showing the variation trend in the total number of genes at each importance cutoff. The knee point was marked as a red dot. **b,** Heatmap showing the expression of selected panel genes in stratified TME groups. The color indicates the z-score-scaled gene expression. Bar plot at the top showing the importance of genes from the random forest model. **c,** Forest plots illustrating the association between the TME-based gene panel and ICB responses in bulk RNA-seq datasets. Coefficients are calculated using a logistic regression model, and the combined coefficient is calculated by random-effects meta-analysis, which is highlighted in red. **d,** Barplot showing the area under the receiver operating characteristic curve (AUC) of our gene panel, other reported gene signatures, and coarse-grained cellular fraction metrics by EPIC when using the XGBoost model. Error bars represent the standard error of the mean. **e,** Bar plots displaying the top 10 dominant cytokines inferred by CytoSig for G05, G07, and G08-G10. **f,** Bar plots showing the differences in average TME group scores between treatment and control groups. Color represents the *P* value. *P* values are calculated by the two-sided Wilcoxon test. ^+^*P*<0.1, \**P* < 0.05, \*\**P* < 0.01, \*\*\**P* < 0.001.

**Fig. S22.**
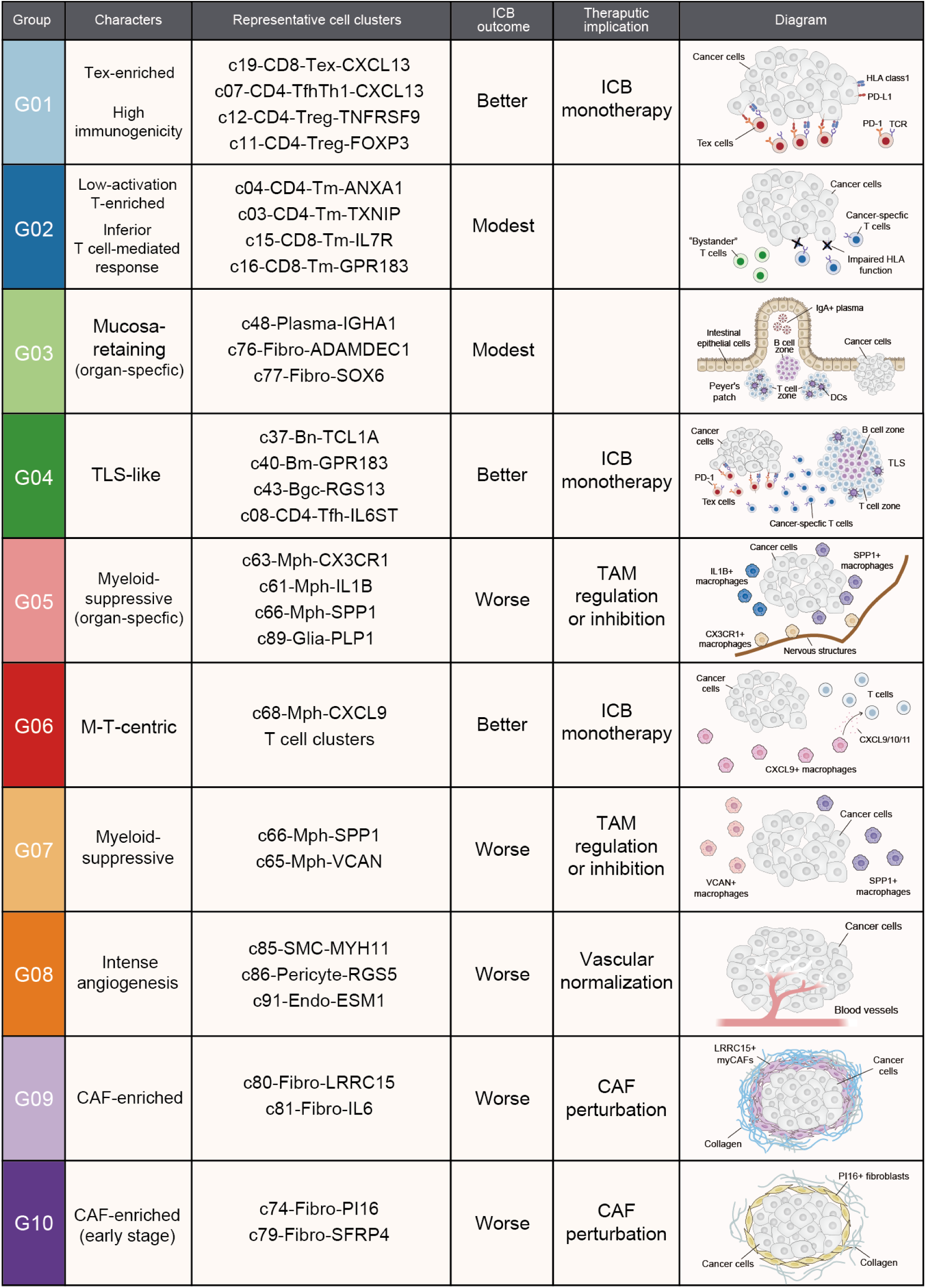
Summary for 10 TME groups.

